# Discovery of Non-Degradative Covalent Molecular Glues for Transcriptional Reprogramming

**DOI:** 10.64898/2025.12.12.694031

**Authors:** Tuong Nghi Duong, Edward Pandji, Qian Shao, Daniel K. Nomura

## Abstract

Transcriptional reprogramming through induced proximity has emerged as a powerful strategy for modulating the expression of oncogenic and tumor-suppressive genes. Inspired by transcriptional reprogramming approaches such as transcriptional/epigenetic chemical inducers of proximity (TCIPs) that link BCL6 inhibitors to transcriptional regulators, we sought to develop covalent ligands that rewire BCL6 proximity to selectively suppress MYC transcriptional output while derepressing BCL6 target loci. Through a chemistry-driven and chemoproteomics-enabled design strategy, we generated a panel of BCL6-based electrophile-bearing hybrid ligands and identified a nondegradative molecular glue, ZD-1-186, that potently suppresses MYC and robustly induces CDKN1A (p21) in diffuse large B-cell lymphoma cells. ZD-1-186 downregulates MYC more effectively than BCL6 inhibitors or degraders, while strongly derepressing canonical BCL6 targets, including p21. Through BCL6 pulldown proteomics, ZD-1-186 induced a selective recruitment of the noncanonical BAF complex subunit BRD9 to BCL6 and covalently modified BRD9 at C288. Pharmacologic inhibition or genetic knockdown of BRD9 attenuated ZD-1-186-mediated MYC suppression and blunted p21 induction. Transcriptomic profiling of ZD-1-186 showed simultaneous derepression of BCL6-repressive loci and suppression of MYC transcriptional programs. These findings demonstrated that ZD-1-186 acted as a transcriptional rewiring glue, recruiting BRD9 to BCL6-repressive loci to activate tumor-suppressive transcription, while also potentially redirecting BCL6 to BRD9-bound oncogenic loci. Overall, our work provides a blueprint for the rational discovery and design of electrophile-enabled, nondegradative molecular glues for targeted transcriptional rewiring.

## Introduction

Targeted reprogramming of transcription through induced proximity is emerging as a promising therapeutic strategy for modulating oncogenic signaling ^1–5^. BCL6 is a BTB zinc finger transcriptional repressor essential for germinal center formation and frequently coopted in diffuse large B cell lymphoma. BCL6 represses tumor suppressor loci, including CDKN1A, TP53 pathway genes, and AP1 family members ^6^. Recent work from Crabtree, Gray, and colleagues introduced transcriptional/epigenetic chemical inducers of proximity (TCIPs) that couple BCL6 inhibitors with ligands for BRD4, CDK9, or EP300 to derepress BCL6 target genes while suppressing oncogenic transcriptional programs ^1–3,6,7^. These heterobifunctional TCIPs revealed that transcriptional outcomes depend not only on blocking BCL6 repressive activity, but also on redirecting BCL6 to new transcriptional environments.

We hypothesized that the modularity of BCL6 ligands could be harnessed to create molecular glues that rewire BCL6 proximity to new transcriptional regulators in a nondegradative manner. To test this, we designed a suite of electrophile-functionalized BCL6 inhibitor hybrids based on the BCL6 inhibitor BI-3812 and the BCL6 degrader BI-3802 scaffolds **(Figure 1a)** ^8,9^. We then implemented a chemoproteomic discovery pipeline to identify constructs that simultaneously derepress tumor suppressor loci and suppress oncogenic MYC transcriptional programs. From this panel, we identified a molecular glue, ZD-1-186, that downregulates MYC expression beyond classical BCL6 inhibitors or degraders, while derepressing BCL6 targets such as CDKN1A. Mechanistically, ZD-1-186 brings together BCL6 with BRD9, a noncanonical BAF complex bromodomain protein implicated in maintaining MYC transcriptional output in specific lymphoma contexts. Through this rewired proximity network, ZD-1-186 recruits BRD9 to BCL6-repression loci to activate transcription, while also potentially bringing BCL6 to BRD9-bound oncogenic loci to suppress MYC-driven programs. Below, we outline the synthesis-driven discovery pipeline, the chemoproteomic mechanism, and the transcriptional consequences of this rewiring.

**Figure 1.**
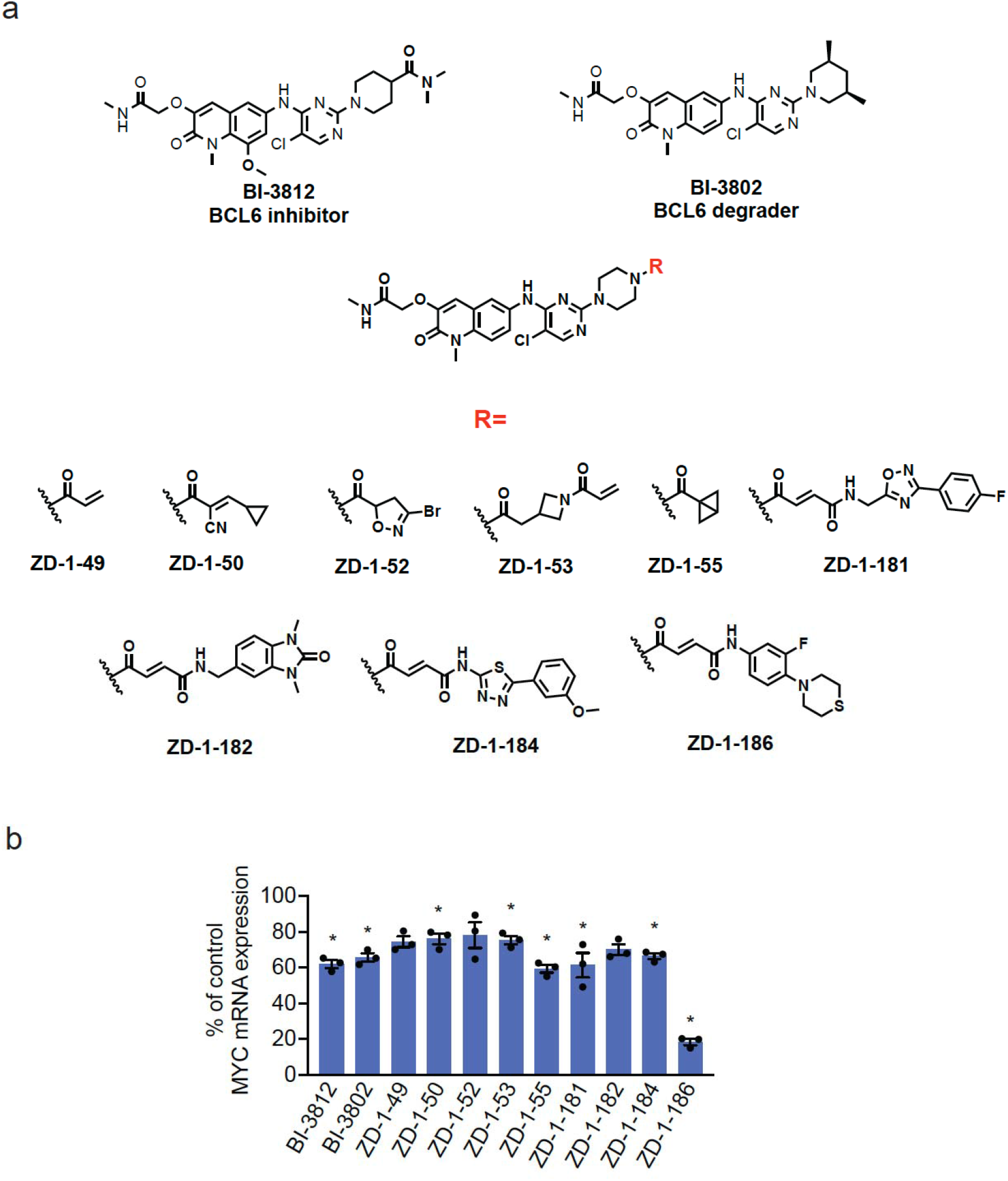
Design and screening of electrophile-appended BCL6 hybrid ligands identify ZD-1-186 as a potent MYC suppressor. **(a)** Structures of BI-3812 (BCL6 inhibitor), BI-3802 (BCL6 degrader), and the electrophile appended hybrid ligands ZD-1-49 through ZD-1-186. Electrophile variants include acrylamides, cyanoacrylates, chloroacetamides, sulfonyl fluorides, and heterocycle-linked handles appended onto the BI-3812 core. **(b)** MYC mRNA expression in HT cells. HT cells were treated with DMSO vehicle or individual compounds (50 μM) for 24 h, after which MYC mRNA levels were assessed by qPCR. ZD-1-186 shows the strongest MYC suppression among the tested analogs, outperforming both BI-3812 and BI-3802. The bar graph shows individual percent replicate values compared to the control from n=3 biologically independent replicates per group, with significance expressed by *p<0.05.

## Results

### Design of Electrophile-Bearing BCL6 Ligands Identifies ZD-1-186 as a MYC Suppressive Hit

We first synthesized a panel of electrophile-appended BCL6 ligands derived from the BI-3812 inhibitor and BI-3802 degrader scaffolds **(Figure 1a)**. These included acrylamides, cyanoacrylates, chloroacetamides, sulfonyl fluorides, and heterocycle-linked electrophiles (**Figure 1a**, structures ZD-1-49 through ZD-1-186). Screening in HT diffuse large B-cell lymphoma (DLBCL) cells revealed that most analogs achieved modest MYC repression, but ZD-1-186 produced a dramatic drop in MYC mRNA levels **(Figure 1b)**. ZD-1-186 was the only hybrid to reduce MYC expression to below 25% of control at screening concentrations. Also, ZD-1-186 only modestly impaired HT cell viability at the highest concentration tested **(Figure S1a)**. This suggested that the electrophile-appended moiety enabled new protein-proximity events that amplified MYC suppression.

### ZD-1-186 Potently Suppresses MYC and Induces CDKN1A Expression in a Dose-Dependent Manner

ZD-1-186 contains the BCL6 inhibitor core fused to a fumaramide-based covalent handle **(Figure 2a)**. Dose-response profiling showed significant suppression of MYC mRNA levels **(Figure 2b)**. We further confirmed a substantial reduction in MYC protein levels and marked induction of CDKN1A (p21) in a dose-dependent manner **(Figure 2c-2d)**. These effects required the electrophile, as the nonreactive analog ZD-2-2 failed to reduce MYC or induce p21 at any concentration tested **(Figure 2e-2f)**. Together, these data indicated that the covalent appendage was essential for the transcriptional rewiring activity of ZD-1-186. We also demonstrated in another BCL6-dependent SU-DHL-4 DLBCL cells that ZD-1-186 similarly reduced MYC expression and strongly induced p21 and another BCL6-repressed target, BIM **(Figure S1b)**.

**Figure 2.**
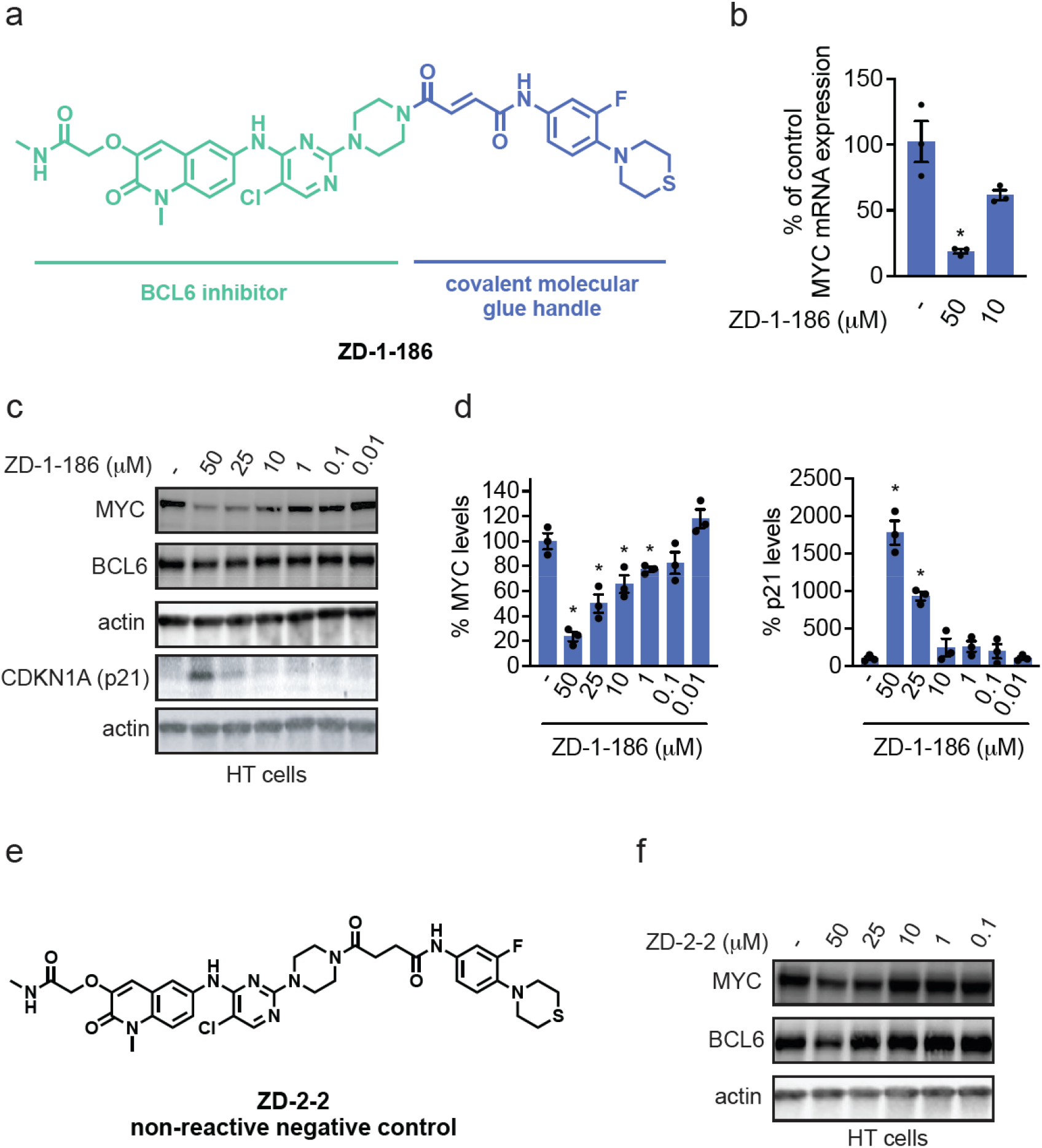
ZD-1-186 potently suppresses MYC and induces CDKN1A in a dose-dependent and electrophile-dependent manner. **(a)** Chemical structure of ZD-1-186 showing the BCL6 inhibitor core and electrophilic covalent handle. **(b)** MYC mRNA levels in HT cells treated with vehicle control or ZD-1-186 for 24 h, after which MYC mRNA levels were assessed by qPCR. **(c)** MYC, BCL6, and CDKN1A (p21) levels in HT cells. HT cells were treated with DMSO vehicle or ZD-1-186 for 16 h and MYC, BCL6, p21, and loading control actin levels were assessed by SDS/PAGE and Western blotting. **(d)** Quantification of MYC and p21 protein levels from **(c). (e)** Chemical structure of the nonreactive negative control analog ZD-2-2 lacking an electrophilic handle. **(f)** MYC, BCL6, and loading control actin levels in HT cells treated with DMSO vehicle or ZD-2-2 for 16 h, assessed by SDS/PAGE and Western blotting. Blots in **(c**,**f)** are representative of n=3 biologically independent replicates per group. The bar graph in **(d)** shows individual percent replicate values compared to the control from n=3 biologically independent replicates per group and average ± sem, with significance expressed by *p<0.05.

### Chemoproteomic Pulldown Identifies BRD9 as a ZD-1-186-Dependent BCL6-Interacting Partner

To uncover the mechanism of transcriptional rewiring, we performed FLAG-BCL6 pulldowns in HEK293T cells treated with ZD-1-186 or vehicle. Proteomic analysis revealed a selective enrichment of BRD9 in the ZD-1-186-treated pulldown **(Figure 3a; Table S1)**, with BRD9 showing the most significant fold increase and p-value among enriched proteins. We further confirmed this interaction by demonstrating robust ZD-1-186-dependent enrichment of BRD9 in pulled-down BCL6 **(Figure 3b-3c)**. These results indicated that ZD-1-186 induced a new induced proximity between BCL6 and BRD9. Because ZD-1-186 contains fumaramide handle, we hypothesized it might covalently react with nucleophilic residues on BRD9. Competition assays of ZD-1-186 against a rhodamine-functionalized cysteine-reactive iodoacetamide (IA-rhodamine) probe demonstrated dose-dependent competition of BRD9 probe-labeling by ZD-1-186 **(Figure S2a)**. Using intact BRD9 protein and ZD-1-186, we identified a covalent adduct on BRD9, C288, via MS/MS peptide mapping **(Figure S2b; Figure 3d)**. Together, these data support a mechanism in which ZD-1-186 binds BCL6 and covalently engages BRD9, creating a proximity-dependent molecular glue.

**Figure 3.**
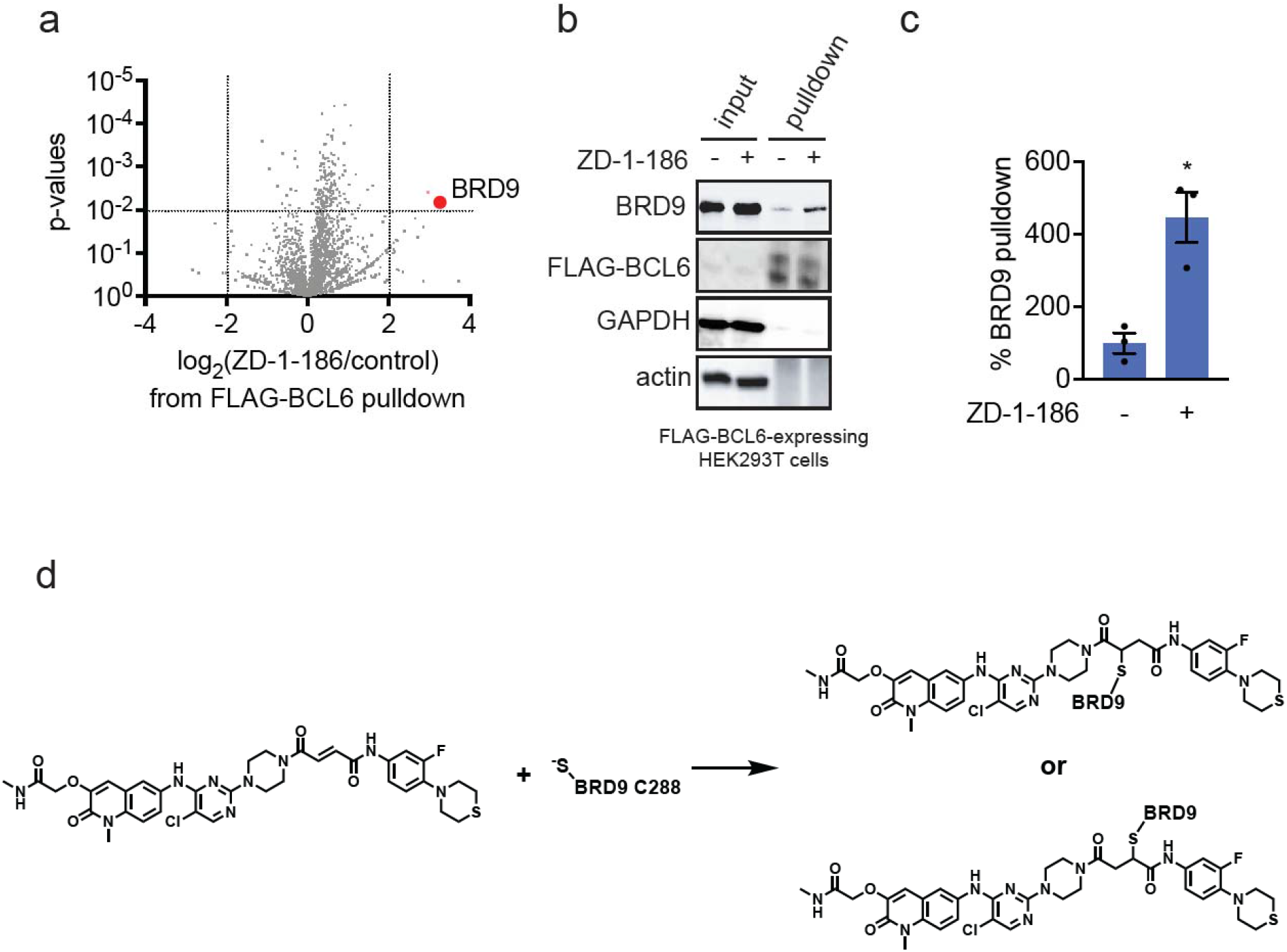
Chemoproteomic profiling reveals BRD9 as a ZD-1-186-dependent BCL6-interacting partner. **(a)** Volcano plot of FLAG-BCL6 pulldown proteomics in FLAG-BCL6-expressing HEK293T cells treated with DMSO vehicle control or ZD-1-186 (50 μM) for 16 h. BRD9 is the most significantly enriched protein upon ZD-1-186 treatment. **(b)** Western blotting validation in FLAG-BCL6 expressing HEK293T cells showing ZD-1-186 dependent enrichment of BRD9 in the BCL6 pulldown. **(c)** Quantification of the experiment described in **(b). (d)** Proposed mechanism showing covalent engagement of BRD9 C288 by ZD-1-186 to generate a stable ternary complex. Proteomics data are provided in **Table S1**. Data in **(a-c)** are from n=3 biologically independent replicates per group. Blot in **(b)** is representative. The bar graph in **(c)** shows individual percent replicate values compared to the control and average ± sem, with significance expressed by *p<0.05.

### BRD9 Functional Perturbation Blunts ZD-1-186 Mediated MYC Suppression and p21 Induction

To test the functional role of BRD9 in this rewired transcriptional program, we evaluated whether BRD9 inhibition antagonized the activity of ZD-1-186. Pre-treatment with the BRD7/9 inhibitor BI-9564, but not with the negative control inhibitor BI-6354, significantly attenuated both MYC suppression and p21 induction in a dose-dependent manner **(Figure 4a-4b)** ^10^. Genetic depletion of BRD9 using shRNA also resulted in similar attenuation of ZD-1-186-mediated MYC suppression **(Figure 4e-4f)**. Importantly, we also demonstrated that neither BRD9 pharmacological inhibition nor genetic ablation lowers MYC levels **(Figure S3a-S3b; Figure 4e-4f)**. These data established BRD9 as a required functional partner for ZD-1-186-driven transcriptional rewiring.

**Figure 4.**
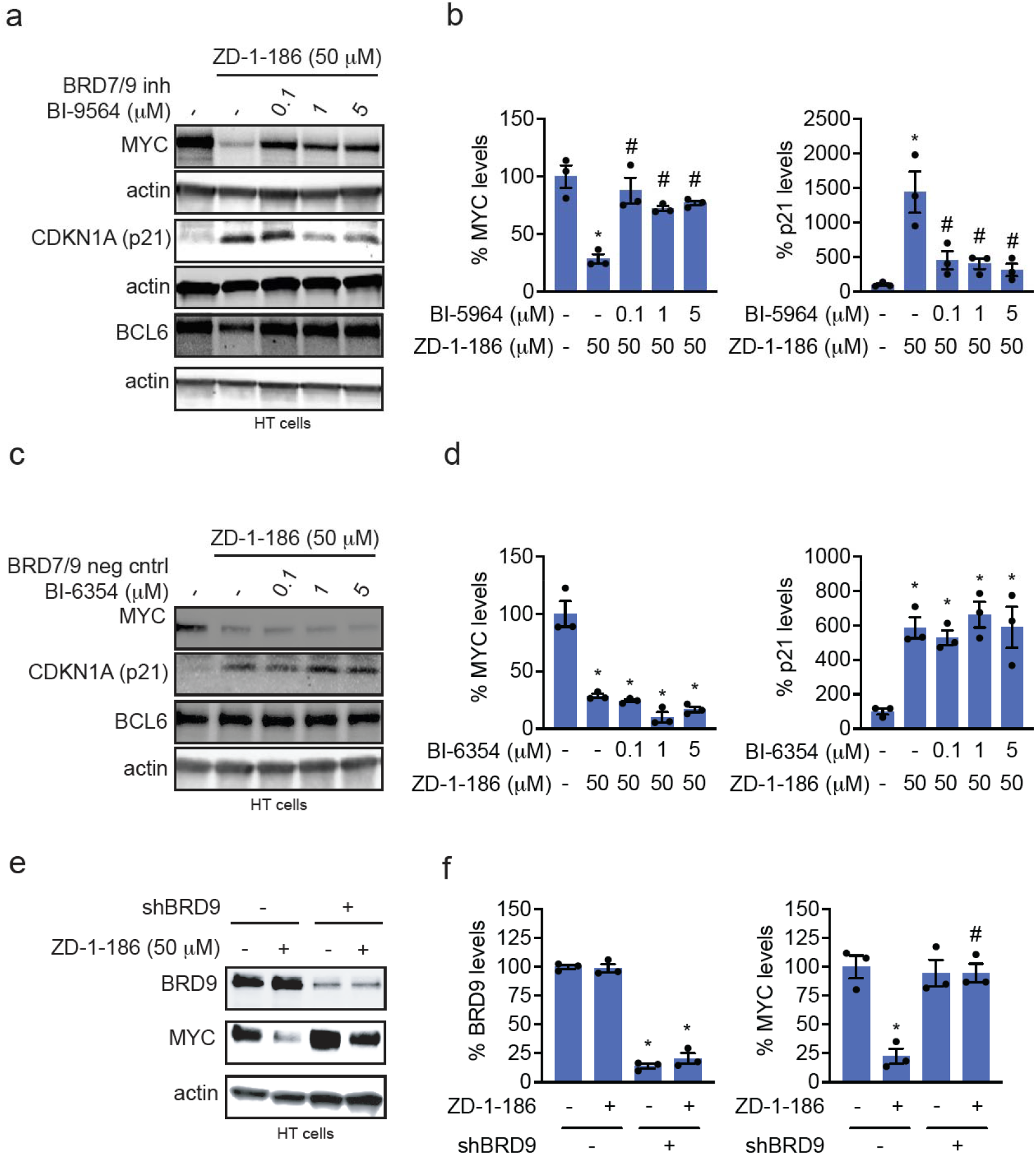
BRD9 is required for ZD-1-186-mediated suppression of MYC and induction of CDKN1A. **(a**,**c)** MYC, p21, BCL6, and loading control actin levels in HT cells co-treated with DMSO vehicle or BRD7/9 inhibitor BI-9564 **(a)** or negative control inhibitor BI-6354 **(c)** alongside treatment of cells with DMSO vehicle or ZD-1-186 for 24 or 16 h, respectively, and assessed by SDS/PAGE and Western blotting. **(b**,**d)** Quantification of MYC and p21 levels from panel **(a**,**c). (e)** BRD9, MYC, and loading control actin levels in control or shBRD9 HT cells treated with DMSO vehicle or ZD-1-186 for 16 h, assessed by SDS/PAGE and Western blotting. **(f)** Quantified BRD9 and MYC levels from panel **(e)**. Blots in **(a**,**c**,**e)** are representative of n=3 biologically independent replicates per group where individual replicates and average ± sem are expressed in bar graphs in **(b**,**d**,**f)**. Significance is expressed as *p<0.05 compared to vehicle treated control cells or #p<0.05 compared to ZD-1-186-treated cells ZD-1-186-treated control cells in **(b**,**d**,**f)**.

### RNA Sequencing Reveals Concurrent Derepression of BCL6 Targets and Suppression of MYC-Driven Transcriptional Programs

RNA sequencing in HT cells treated with ZD-1-186 revealed distinct classes of transcriptional changes. A strong derepression was observed at canonical BCL6-repressive loci, including NR4A family members, AP1 factors, DDIT3, CDKN1B, TP53INP1, and stress-response genes **(Figure 5a; Table S2)**. Simultaneously, MYC-dependent transcriptional programs, including MYC, E2F, oxidative phosphorylation, and biosynthetic pathways, were strongly downregulated **(Figure 5b; Table S2)**. Pathway analysis clearly segregated these signatures into significantly derepressed BCL6 targets and significantly suppressed MYC networks **(Figure 5c; Table S2)**. These results support a model in which ZD-1-186 redirected BRD9 to BCL6-repressive loci to activate transcription, while either sequestering BRD9 away from MYC regulatory elements or diverting BCL6 to BRD9-bound MYC regulatory elements to suppress MYC transcription.

**Figure 5.**
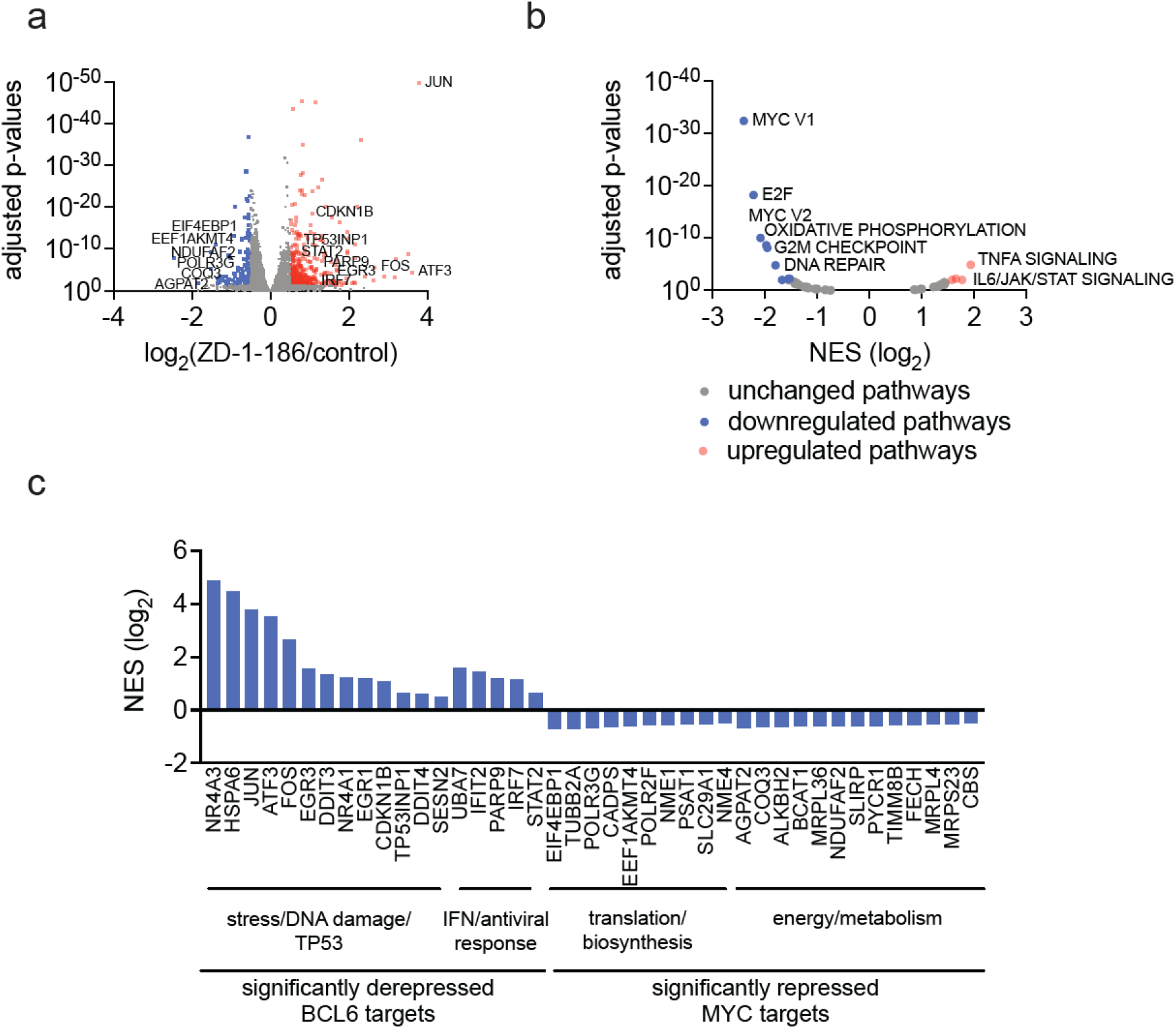
Transcriptional profiling reveals ZD-1-186-driven derepression of BCL6 targets and suppression of MYC transcriptional programs. **(a)** Volcano plot of genes significantly altered by ZD-1-186 in HT cells with representative BCL6 and MYC target genes annotated from RNA sequencing of HT cells treated with DMSO vehicle or ZD-1-186 (50 μM) for 20 h. Canonical BCL6 repressed genes, including NR4A family, AP1 members, DDIT3, CDKN1B, and TP53 pathway genes, are strongly derepressed. **(b)** Gene set enrichment analysis of hallmark pathways from **(a)** showing significant downregulation of MYC, E2F, oxidative phosphorylation, and biosynthetic pathways upon ZD-1-186 treatment. **(c)** Normalized enrichment scores of representative upregulated and downregulated gene sets from **(a)**, showing clear separation between derepressed BCL6 targets and repressed MYC-dependent pathways. RNA sequencing data are from n=3 biologically independent replicates per group and are provided in **Table S2**.

## Discussion

Transcriptional rewiring through induced proximity has recently emerged as a powerful strategy for redirecting gene regulatory circuits that have historically been inaccessible to small molecules. Foundational work from Crabtree, Gray, and colleagues demonstrated that heterobifunctional transcriptional chemical inducers of proximity (TCIPs) can redirect BCL6 to specific transcriptional machinery, thereby modulating gene expression outcomes ^1–4,7^. We also recently reported a heterobifunctional targeted transcriptional repression approach that induces the proximity of a transcriptional repression complex to specific transcription factors, thereby repressing transcription at these loci (Targeted Transcriptional Regulation via Active Control of Epigenetic Reprogramming, or TRACERs) ^5^. The Gibson lab also recently demonstrated transcriptional rewiring of p53 tumor-suppressive loci using targeted protein localization approaches ^11^. These studies have established the conceptual framework that transcription factors can be functionally reprogrammed not only by inhibiting their endogenous activity but by relocating them to new regulatory contexts. Building on this foundation, our work extends these principles to the design of nondegradative, covalency-enabled molecular glues that rewire transcriptional networks via endogenous protein surfaces, without requiring bifunctional architectures.

Our data, as observed with the heterobifunctional TCIPs^2^, reveal that ZD-1-186 induces a dual rewiring mechanism in which BRD9 is redirected to BCL6-repressive loci to activate tumor-suppressive transcription, and either BRD9 is sequestered away from MYC regulatory elements, or BCL6 is redirected to BRD9-bound oncogenic loci. This bidirectional rewiring simultaneously suppresses MYC-driven transcriptional programs and derepresses canonical BCL6 targets, including p21, TP53 pathway genes, AP1 family members, and NR4A factors. Importantly, this outcome is not achieved by BCL6 inhibition or degradation alone but requires the formation of a ZD-1-186-dependent BCL6-BRD9 proximity network, supported by proteomics, covalent engagement of BRD9 C288, pharmacologic rescue, and genetic loss of BRD9 function.

One may reasonably ask whether the activity of ZD-1-186 reflects generalized BAF perturbation rather than a specific BCL6-BRD9 rewiring mechanism. Several observations argue against this. First, BRD9 engagement requires a covalent electrophile; the nonreactive analog ZD-2-2 fails to recapitulate MYC suppression or p21 induction. Second, pharmacologic inhibition of BRD9 with BI-9564 selectively attenuates the effects of ZD-1-186, whereas the BRD7 selective negative control BI-6354 does not. Third, BRD9 knockdown blunts MYC suppression even in the continued presence of ZD-1-186. Fourth, neither BRD7/9 pharmacological inhibition nor BRD9 genetic knockdown alone downregulates MYC levels. Together, these data indicate that BRD9 is not a bystander but an obligate cofactor in the transcriptional response elicited by ZD-1-186. Another concern is whether the observed transcriptional changes arise from indirect stress responses rather than targeted rewiring of BCL6-mediated regulatory circuits. While ZD-1-186 upregulates stress-responsive genes, these signatures align closely with known BCL6-repressive networks and do not resemble nonspecific cytotoxic transcriptional programs. Furthermore, MYC suppression does not correlate with a loss of cell viability at the tested time points. Instead, the transcriptomic data are consistent with a rewired promoter-occupancy model, in which BRD9 and BCL6 are redistributed to each other’s regulatory environments. Although direct chromatin occupancy studies such as CUT and RUN or CUT and Tag could further substantiate this model, the combined proteomic, covalent-labelling, pharmacologic, and genetic data provide a coherent mechanistic basis for transcriptional rewiring.

Finally, the question of generalizability is central to evaluating the impact of any new molecular glue platform. We and others have previously demonstrated that electrophile-functionalized ligands can be used to identify permissive covalent degradative handles and E3 ligase pairs for rationally designing molecular glue degraders ^12–18^. Our study provides proof of principle that carefully tuned electrophilic appendages can convert traditional transcription factor inhibitors into molecular glues that function not by degrading or inhibiting proteins, but by redirecting protein surfaces into new regulatory complexes. This approach complements, rather than replaces, heterobifunctional TCIP architectures and expands the design space for transcriptional reprogramming. Of future interest will be whether this covalent handle can be transplanted onto other transcription factor-targeting ligands to modulate the transcriptional programming of other targets through BRD9 recruitment. Of additional interest will be whether this electrophile-functionalized ligand approach can also be used more generally to identify additional covalent handles and epigenetic effector pairs for molecular glue TCIPs and targeted transcriptional repressors (TRACERs).

In summary, our work exemplifies a new strategy for nondegradative molecular glue design that uses chemoproteomic and electrophile-driven principles to rewire oncogenic transcriptional circuits. Building on the foundational insights from TCIP technology, our work establishes a framework for rationally engineering glues that modulate gene expression not by destroying or blocking transcription factors but by redirecting them into new functional proximities. As molecular glue discovery shifts increasingly toward targeted transcriptional control, this chemistry-first, mechanism-based approach enables tuning of transcriptional states with precision.

## Acknowledgment

We thank the members of the Nomura Research Group and Novartis BioMedical Research for their critical review of the manuscript. This work was also supported by Novartis Biomedical Research, the National Science Foundation Molecular Foundations for Biotechnology (MFB) grant (2127788), the UC Berkeley Molecular Therapeutics Initiative (MTI), the Mark Foundation for Cancer Research ASPIRE Award, the National Institutes of Health (R35CA263814, R01CA240981), and the Bakar Award. We also thank Hasan, Lund, and the UC Berkeley NMR facility in the College of Chemistry (CoC-NMR) for spectroscopic assistance. Instruments in the College of Chemistry NMR facility are partly supported by NIH S10OD024998. Boehringer Ingelheim kindly provided BI-6354 via its open innovation platform opnMe, available at https://opnme.com.

## Author Contributions

TND, QS, and DKN conceived of the project. TND, EP, QS, and DKN designed, performed, and analyzed experiments. TND, QS, and DKN wrote the paper.

