## Supporting Information for "Discovery of Non-Degradative Covalent Molecular Glues for Transcriptional Reprogramming"

[1] Departments of Chemistry and Molecular and Cell Biology. University of California, Berkeley, Berkeley, CA 94720 USA

[2] Innovative Genomics Institute, Berkeley, CA 94720 USA

[3] Molecular Therapeutics Initiative, Berkeley, CA 94720 USA

[4] Novartis-Berkeley Translational Chemical Biology Institute, Berkeley, CA 94720 USA

**Supporting Table Legends**

**Table S1. FLAG-BCL6 pulldown proteomics**. FLAG-pulldown TMT-based quantitative proteomics in FLAG-BCL6-expressing HEK293T cells treated with DMSO vehicle control or ZD-1-186 (50 μM) for 16 h.

**Table S2. Transcriptional profiling reveals ZD-1-186-driven derepression of BCL6 targets and suppression of MYC transcriptional programs.** HT cells treated with DMSO vehicle or ZD-1-186 (50 μM) for 20 h, RNA was extracted and subjected to RNA sequencing. Tab 1 shows total RNA sequencing data, ratio of treated versus control and statistical significance expressed as adjusted p-values. Tab 2 shows gene set enrichment analysis of hallmark pathways. Tab 3 shows normalized enrichment scores for representative upregulated and downregulated genes, clearly separating derepressed BCL6 targets from repressed MYC-dependent pathways.

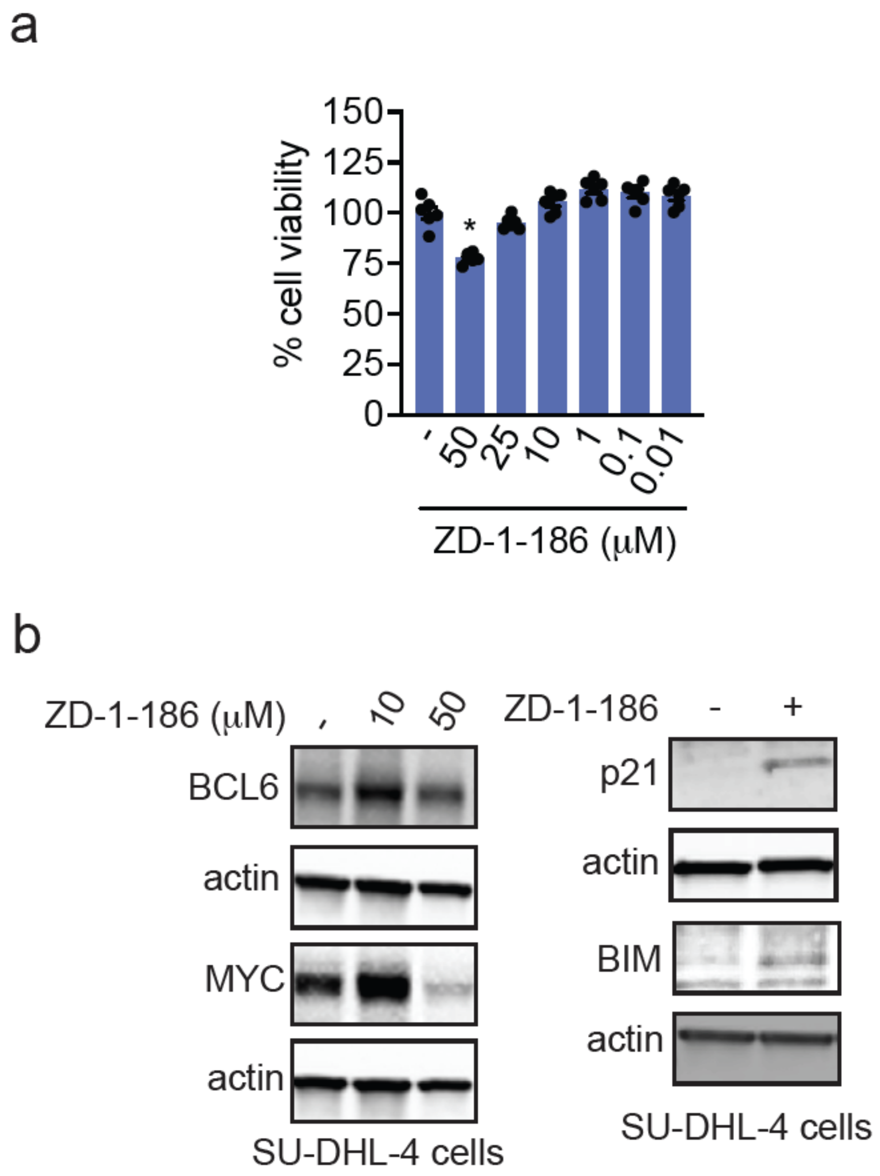


**Figure S1. ZD-1-186 activity in DLBCL cell lines. (a)** ZD-1-186 effects on cell viability. HT cells were treated with DMSO vehicle or ZD-1-186 for 16 h, after which cell viability was assessed. **(b)** BCL6, MYC, p21, BIM, and loading control actin levels in SU-DHL-4 cells treated with DMSO vehicle or ZD-1-186 (designated concentration or 50 μM) for 24 h, assessed by SDS/PAGE and Western blotting. Blots are representative of n=3 biologically independent replicates per group. Data in **(a)** shows individual replicate values and average ± sem from n=6 biologically independent replicates per group.

a

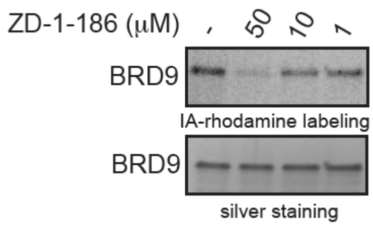


b

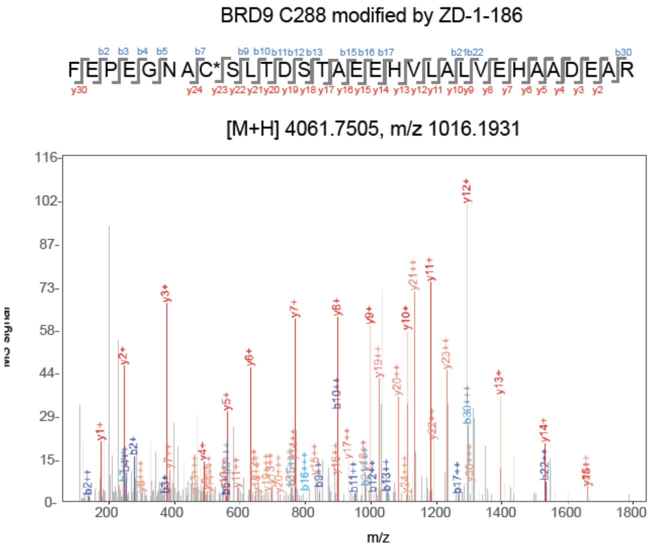


| b+ | b2+ | b3+ | # | Seq | # | y+ | y2+ | y3+ |
|---|---|---|---|---|---|---|---|---|
| 148.0757 | 74.5415 | 50.0301 | 1 | F | 31 | | | |
| 277.1183 | 139.0628 | 93.0443 | 2 | E | 30 | 3914.6820 | 1957.8447 | 1305.5655 |
| 374.1710 | 187.5892 | 125.3952 | 3 | P | 29 | 3785.6394 | 1893.3234 | 1262.5513 |
| 503.2136 | 252.1105 | 168.4094 | 4 | E | 28 | 3688.5867 | 1844.7970 | 1230.2004 |
| 560.2351 | 280.6212 | 187.4166 | 5 | G | 27 | 3559.5441 | 1780.2757 | 1187.1862 |
| 674.2780 | 337.6427 | 225.4309 | 6 | N | 26 | 3502.5226 | 1751.7649 | 1168.1791 |
| 745.3151 | 373.1612 | 249.1099 | 7 | A | 25 | 3388.4797 | 1694.7435 | 1130.1647 |
| 1598.5627 | 799.7850 | 533.5257 | 8 | C | 24 | 3317.4426 | 1659.2249 | 1106.4857 |
| 1685.5947 | 843.3010 | 562.5364 | 9 | S | 23 | 2464.1950 | 1232.6012 | 822.0699 |
| 1798.6788 | 899.8430 | 600.2311 | 10 | L | 22 | 2377.1630 | 1189.0851 | 793.0592 |
| 1899.7265 | 950.3669 | 633.9137 | 11 | T | 21 | 2264.0789 | 1132.5431 | 755.3645 |
| 2014.7534 | 1007.8803 | 672.2560 | 12 | D | 20 | 2163.0313 | 1082.0193 | 721.6819 |
| 2101.7854 | 1051.3964 | 701.2667 | 13 | S | 19 | 2048.0043 | 1024.5058 | 683.3396 |
| 2202.8331 | 1101.9202 | 734.9492 | 14 | T | 18 | 1960.9723 | 980.9898 | 654.3289 |
| 2273.8702 | 1137.4388 | 758.6283 | 15 | A | 17 | 1859.9246 | 930.4659 | 620.6464 |
| 2402.9128 | 1201.9600 | 801.6425 | 16 | E | 16 | 1788.8875 | 894.9474 | 596.9674 |
| 2531.9554 | 1266.4813 | 844.6567 | 17 | E | 15 | 1659.8449 | 830.4261 | 553.9532 |
| 2669.0143 | 1335.0108 | 890.3430 | 18 | H | 14 | 1530.8023 | 765.9048 | 510.9390 |
| 2768.0827 | 1384.5450 | 923.3658 | 19 | V | 13 | 1393.7434 | 697.3753 | 465.2527 |
| 2881.1668 | 1441.0870 | 961.0605 | 20 | L | 12 | 1294.6750 | 647.8411 | 432.2298 |
| 2952.2039 | 1476.6056 | 984.7395 | 21 | A | 11 | 1181.5909 | 591.2991 | 394.5352 |
| 3065.2880 | 1533.1476 | 1022.4342 | 22 | L | 10 | 1110.5538 | 555.7805 | 370.8561 |
| 3164.3564 | 1582.6818 | 1055.4570 | 23 | V | 9 | 997.4697 | 499.2385 | 333.1614 |
| 3293.3990 | 1647.2031 | 1098.4712 | 24 | E | 8 | 898.4013 | 449.7043 | 300.1386 |
| 3430.4579 | 1715.7326 | 1144.1575 | 25 | H | 7 | 769.3587 | 385.1830 | 257.1244 |
| 3501.4950 | 1751.2511 | 1167.8365 | 26 | A | 6 | 632.2998 | 316.6536 | 211.4381 |
| 3572.5321 | 1786.7697 | 1191.5156 | 27 | A | 5 | 561.2627 | 281.1350 | 187.7591 |
| 3687.5591 | 1844.2832 | 1229.8579 | 28 | D | 4 | 490.2256 | 245.6164 | 164.0801 |
| 3816.6017 | 1908.8045 | 1272.8721 | 29 | E | 3 | 375.1987 | 188.1030 | 125.7377 |
| 3887.6388 | 1944.3230 | 1296.5511 | 30 | A | 2 | 246.1561 | 123.5817 | 82.7235 |
| | | | 31 | R | 1 | 175.1190 | 88.0631 | 59.0445 |

Static Modifications:
**C: 57.02146**

Variable Modifications:
**C: 693.2169**

**Figure S2. ZD-1-186 covalently modifies BRD9 at C288. (a)** IA-rhodamine competition assay showing dose-dependent reduction in BRD9 labeling in the presence of ZD-1-186. BRD9 pure protein was pre-incubated with DMSO vehicle or ZD-1-186 for 2h prior to labeling of the protein with IA-rhodamine (250 nM) for 1 h. IA-rhodamine labeling was assessed by SDS/PAGE and in-gel fluorescence and loading was assessed by silver staining. **(b)** MS/MS spectrum and peptide mapping of BRD9, identifying a covalent adduct at cysteine 288 formed by ZD-1-186. BRD9 pure protein was treated with ZD-1-186 (50 μM) for 1h after which the protein was digested with trypsin and tryptic digests were analyzed by LC-MS/MS to identify the ZD-1-186-modified tryptic peptide. Shown are MS/MS spectra and analysis and the modified cysteine. Data are representative of n=3 biologically independent replicates per group.

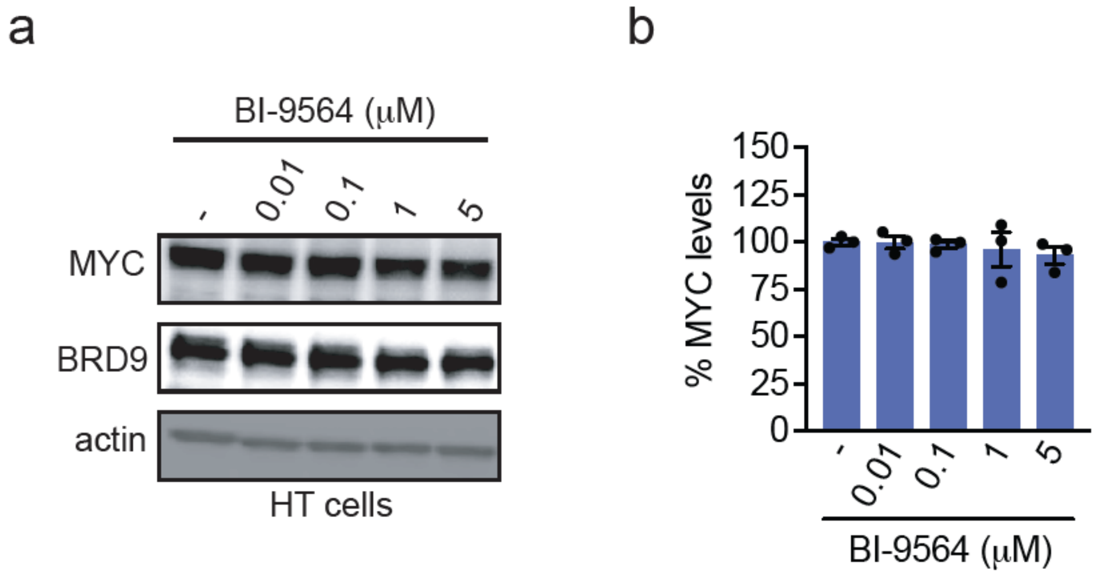


**Figure S3. BRD7/9 inhibitor BI-9564 does not downregulate MYC. (a, b)** HT cells were treated with DMSO vehicle or BI-9564 for 24 h, after which MYC, BRD9, and loading control actin levels were assessed by SDS/PAGE and Western blotting **(a)** and quantified in **(b).** Blot shown in **(a)** is representative of n=3 biologically independent replicates. Bar graph in **(b)** shows individual replicate values and average ± sem.

### Materials and Methods

#### Cell Culture

HT cells were obtained from American Tissue Culture Collection (ATCC). Lymphoma cells were grown in RPMI-1640 medium supplemented with 10% (v/v) fetal bovine serum (FBS). HEK293T cells were obtained from the UC Berkeley Cell Culture Facility and grown in Dulbecco's Modified Eagle Medium (DMEM) supplemented with 10% (v/v) FBS and in incubators. HT BRD9 knockout cells were obtained from Signo Biological and cultured in RPMI-1640 medium supplemented with 10% (FBS). All cell lines were incubated at 37°C with 5% $CO_2$.

#### Western Blotting

Cells were plated at 1 M $ml^{-1}$ and treated with drug at indicated timepoints and doses. Cells were harvested in cold PBS buffer and pelleted by centrifugation (1,300 g, 5 min, 4°C). Pelleted cells were resuspended in a mixture of RIPA lysis buffer (Thermo Fisher) and Halt Protease and Phosphatase Inhibitor Cocktail (Thermo Fisher, 78440) and incubated for 30 minutes on ice. Cell lysates were centrifuged (20,000 g, 10 min, 4°C). The supernatant was collected to pre-chilled microcentrifuge tubes. Protein concentration was measured by BCA assay and normalized to equal concentration. SDS-PAGE analysis was carried out in precast 4-20% Criterion TGX Gels (Bio-Rad). Proteins were transferred to nitrocellulose membrane using the Trans-Blot Turbo transfer system (Bio-Rad). Membranes were blocked with 5% BSA in Tris-buffered saline containing Tween 20 (TBS-T) solution for 1 h at RT and subsequently probed with primary antibody diluted solutions overnight at 4°C. Membranes were washed with TBS-T three times before incubating with IR-680 and IR-800 secondary antibodies at 1:5,000 dilution in 5% BSA in TBS-T for 1 h at RT. Membranes were stripped using ReBlot Plus Strong Antibody Stripping Solution (EMD Millipore), when additional primary antibody incubations were performed. Antibodies used for immunoblots were: BCL6 (1:1000 v/v, Cell Signaling, D65C10), MYC (1:1000 v/v, Cell Signaling, D84C12), p21 (1:500 v/v, Abcam, EPR3993), DYKDDDDK Tag (1:1000 v/v, Cell Signaling, D6W5B), Actin (1:20000 v/v, Protein Tech, 66009-1-Ig), GAPDH (1:50000 v/v, Protein tech, 60004-1-Ig). Odyssey (Li-Cor) scanner was used for blot imaging.

#### Cell Viability Assay

Cells were plated at 0.1M $ml^{-1}$ in 96-well white plates then treated with drug at indicated timepoints and dose. Cell viability assay was performed using CellTiter-Glo 2.0 reagent (Promega, G9241) according to manufacturer's protocol. Luminescent signals were measured using the Tecan Spark Plate reader (30086376).

#### RT-qPCR

Cells were plated at 1M $ml^{-1}$ and treated with drug at indicated timepoints and doses. Cells were harvested in cold PBS buffer. Total RNA was extracted using Monarch Total RNA Miniprep Kit (NEB, T2010S) Kit (NEB, T2010S). cDNA was prepared using Luna Universal One-Step RT-qPCR Kit (NEB, E3005S). Gene expression was measured from cDNA using CFX Connect Real-Time PCR Detection System (BioRad). Relative gene expression was normalized to housekeeping genes (GAPDH, β-actin) and calculated using comparative Ct method. Primer sequences were the following:

*BCL6* (forward): CATGCAGAGATGTGCCTCCACA
*BCL6* (reverse): TCAGAGAAGCGGCAGTCACACT
*c-MYC* (forward): CCTGGTGCTCCATGAGGAGAC
*c-MYC* (reverse): CAGACTCTGACCTTTTGCCAGG
*p-21* (forward): AGGTGGACCTGGAGACTCTCAG
*p-21* (reverse): TCCTCTTGGAGAAGATCAGCCG
*GAPDH* (forward): GTCTCCTCTGACTTCAACAGCG
*GAPDH* (reverse): ACCACCCTGTTGCTGTAGCCAA
*β-actin* (forward): CACCATTGGCAATGAGCGGTTC
*β-actin* (reverse): AGGTCTTTGCGGATGTCCACGT

#### RNA Sequencing

Total RNA was extracted as described above. RNA sequencing library preparation and sequencing were carried out by the QB3-Berkeley Genomics Facility (QB3 Genomics, UC Berkeley, Berkeley, CA, RRID:SCR_022170). RNA integrity and poly-dT–enriched mRNA quality was assessed using an Agilent 2100 Bioanalyzer. Libraries were generated with the KAPA mRNA HyperPrep Kit (Roche KK8581). Truncated universal stub adapters were ligated to cDNA fragments and extended into full-length Illumina libraries through 10 cycles of PCR using unique dual-index primers. Library quality was evaluated on an AATI (now Agilent) Fragment Analyzer before being transferred to the Vincent J. Coates Genomics Sequencing Laboratory (GSL), another QB3-Berkeley Core Research Facility at UC Berkeley.

**RNA Sequencing Analysis**

Differential expression analysis Samples were sequenced on one lane of a NovaSeq X Plus generating paired-end 150bp reads to a median depth of 59.3 million reads per sample (minimum 46.3 million reads). After demultiplexing, raw FASTQ files were aligned and quantified using Kallisto[1] and filtered for >10 transcripts per million (tpm) for all replicates of at least one condition followed by differential gene expression analysis using DESeq2[2] and a threshold of |log2(FoldChange)| > 1 and an adjusted $p$-value < 0.05 was applied to identify differentially expressed genes. fGSEA analysis[3] was performed to identify significantly regulated MSigDB Hallmark gene sets[4].

**Gel-based ABPP**

Recombinant BRD9 (Origene, TP760173) (0.25 μg/sample) was pretreated with either DMSO or covalent ligand at 37°C for 2 h in 25 μL of PBS. Samples were treated with 0.25 μM IA-Rhodamine (Setareh Biotech) at RT for 1 h in the dark. The reaction was quenched by adding 4X reducing Laemmli SDS loading buffer (Alfa Aesar). Samples were boiled at 95°C for 5 min and separated on precast 4-20% Criterion TGX Gels (Bio-Rad). Probe-labeled proteins were analyzed using in-gel fluorescence and visualized on Chemidoc MP (Bio-Rad). Imaged gels were stained using Pierce Silver Stain Kit (Thermo Fisher, 24612) according to the manufacturer's protocol.

**BRD9 Lentiviral Knockdown Studies in HT cells**

MISSION shRNA lentiviral construct, pMD2.G (Addgene, 12259), and psPAX2 (Addgene, 12260) were transfected into HEK293T cells using Lipofectamine 2000 (Thermo Fisher, 11668027). Transfected cells were incubated for 48 h at 37°C with 5% $CO_2$. After 48 h, the virus-containing medium was collected, filtered, supplemented with 1:1000 dilution of Polybrene (Sigma-Aldrich, TR-1003-G), and used to infect target cells (HT, ATCC, CRL-2260). The infected cells were incubated for 48 h at 37°C with 5% $CO_2$ and subsequently selected with puromycin. MISSION pLKO.1-puro non-mammalian shRNA Control (Sigma-Aldrich, SHC016) was used as the control shRNA. shRNA sequences were the following:
shBRD9 (Sigma-Aldrich, TRCN0000127634): CGAGTAGAGAAGTTATCAGCT
shBRD9 (Sigma-Aldrich, TRCN0000131081): GAGAGCACACCTATTCAGCAA
shBRD9 (Sigma-Aldrich, TRCN0000236473): CTCCTGGATATTCAATGATAA

**Mapping of ZD-1186 Site of Modification on BRD9 by LC-MS/MS**

Recombinant BRD9 protein (20 μg, Origene, TP760173) was diluted in 100 μL of PBS and pretreated with 50 μM ZD-1186 for 1 h at RT. The protein was precipitated by adding 25 μL of cold acetone and incubated for 1 h at -20°C. The sample was centrifuged (20,000 g, 10 min, 4°C). The supernatant was carefully removed. The pelleted sample was washed with 200 μL of ice-cold 0.01 M HCl/90% acetone solution, followed by centrifugation (20,000 g, 5 min, 4°C); this step was repeated three times. The sample was resuspended and vortexed in 30 μL of 8M Urea in PBS and 30 μL of ProteaseMax surfactant (20 μg $ml^{-1}$ in 100 mM ammonium bicarbonate, Promega, V2071). 40 μL of 100 mM ammonium bicarbonate was added to reach a final volume of 100 μL. The sample was reduced with 10 μL of TCEP (10 mM final concentration) for 30 min at 60°C and subsequently alkylated with 10 μL of iodoacetamide (12.5 mM final concentration) for 30 min at 37°C. The sample was diluted with 120 μL of PBS and added 1.2 μL of ProteaseMax surfactant (0.1 mg $ml^{-1}$ in 100 mM ammonium bicarbonate) along with 10 μL of sequencing grade trypsin (0.5 mg $ml^{-1}$ in 50mM ammonium bicarbonate), followed by overnight incubation at 37°C. The next day, the sample was acidified with 12 μL of formic acid (5% final concentration), dried using a vacuum concentrator at 30°C, and fractionated using high pH reverse-phase peptide fractionation kit (Thermo Fisher, 84868) according to the manufacturer's protocol.

**Generation of FLAG-tagged BCL6 overexpression in HEK293T cells**

Lentiviral particles expressing FLAG-BCL6 (Origene, RC219007L3V) were transduced into HEK293T cells with the addition of polybrene (8 μg $ml^{-1}$ final concentration) according to manufacturer's protocol. Transduced cells were incubated for at 37°C with 5% $CO_2$. The next day, medium containing lentiviral particles was removed and replaced with pre-warmed complete culture medium. After 24 h of resting, cell medium was replaced and selected with pre-warmed complete culture medium supplemented with puromycin (2.5 μg $ml^{-1}$ final concentration).

**Anti-FLAG Pulldown**

HEK293T cells with stable expression of FLAG-BCL6 were treated with DMSO vehicle or 50 μM ZD-1-186 for 16 h. Cells were collected, washed twice with PBS, and pelleted (1,300 g, 5 min, 4°C). Pelleted cells were resuspended in IP Lysis buffer supplemented with protease inhibitor cocktail (Pierce, A32955) and incubated for 30 min at 4°C. Cell lysates were centrifuged (20,000g, 10 min, 4°C). The supernatant was transferred to new conical tubes and incubated with Anti-DYKDDDDK G1 affinity resin (GenScript) for 3 h at RT. Resins were washed twice with PBS and water. The pulldown samples were subsequently prepared for western blot as described above and proteomic analysis as described under Proteomic analysis of pulldown samples.

**Proteomic Analysis of Pulldown Samples**

The samples were diluted in 350 μL urea (6 M in PBS), treated with DTT (25 μL, 30 mg $ml^{-1}$ in water), and incubated at 65°C for 30 min. The samples were reacted with iodoacetamide (25 μL, 74 mg $ml^{-1}$ in water) and incubated with agitation at 37°C for 30 min. The samples were diluted with 720 μL of PBS, added $CaCl_2$ (2 μL, 100 mM, 14.7 mg $ml^{-1}$), and digested with trypsin (4 μL, 0.5 mg $ml^{-1}$) overnight at 37°C. The samples were acidified with formic acid (5% final concentration). The sample supernatants were washed and eluted via C18 purification (Water, WAT051910) following the manufacture's protocol. The eluent was dried using a vacuum concentrator at 30°C. The digested peptides were labeled using commercially available TMT sixplex tags (Thermo Fisher, V5111). After labeling, 35 μg of each labeled sample was combined and dried using a vacuum concentrator at 30°C. The dried samples were redissolved in 300 μL of 0.1% TFA in water and further fractionated using high-pH reverse-phase peptide fractionation kits (Thermo Fisher, 84868) following the manufacturer's protocol. The dried fractions were resuspended in 100 μL of 0.1% formic acid in water for LC-MS/MS analysis.

**Mass Spectrometry Analysis**

Mass spectrometry analysis was performed on an Orbitrap Eclipse Tribrid Mass Spectrometer with a High Field Asymmetric Waveform Ion Mobility (FAIMS Pro) Interface (Thermo Scientific) with an UltiMate 3000 Nano Flow Rapid Separation LCnano System (Thermo Scientific). Offline fractionated samples (5 μL aliquot of 15 μL sample) were injected via an autosampler (Thermo Scientific) onto a 5 μL sample loop, which was subsequently eluted onto an Acclaim PepMap 100 C18 HPLC column (75 μm × 50 cm, nanoViper). The peptides were separated at a flow rate of 0.3 μL/min using the following gradient: 2% buffer B (100% acetonitrile with 0.1% formic acid) in buffer A (95:5 water/acetonitrile, 0.1% formic acid) for 5 min, followed by a gradient from 2 to 40% buffer B from 5 to 159 min, 40 to 95% buffer B from 159 to 160 min, held at 95% B from 160 to 179 min, 95% to 2% buffer B from 179 to 180 min, and then 2% buffer B from 180 to 200 min. The voltage applied to the nano-LC electrospray ionization source was 2.1 kV. Data were acquired through an MS1 master scan (Orbitrap analysis, resolution 120,000, 400−1800 m/z, RF lens 30%, heated capillary temperature 250 °C) with dynamic exclusion (repeat count 1, duration 60 s). Datadependent data acquisition comprised a full MS1 scan, followed by sequential MS2 scans based on 2 s cycle times. FAIMS compensation voltages (CVs) of −35, −45, and −55 were applied. MS2 analysis consisted of a quadrupole isolation window of 0.7 m/z of the precursor ion followed by a higher energy collision dissociation (HCD) energy of 38% with an orbitrap resolution of 50,000. Quantitative TMT-based proteomic analysis was performed, as previously described using a Thermo Eclipse with FAIMS LC-MS/MS.32 Acquired MS data was processed using ProLuCID search methodology in IP2 v.3-v.5 (Integrated Proteomics Applications, Inc.).33 Trypsin cleavage specificity (cleavage at K, R, except if followed by P) allowed for up to 2 missed cleavages. Carbamidomethylation of cysteine was set as a fixed modification, and methionine oxidation and TMT modification of N-terminus, and lysine residues were set as variable modifications. Reporter ion ratio calculations were performed using summed abundances with the most confident centroid selected from the 20 ppm window. Only peptide-to-spectrum matches that are unique to a given identified protein within the total data set are considered for protein quantitation. High-confidence protein identifications were reported with a <1% false discovery rate (FDR) cutoff. Differential abundance significance was estimated using ANOVA with Benjamini− Hochberg correction to determine the p-values.

**Synthetic Methods and Characterization**

All chemical reactions were performed under a nitrogen atmosphere with dry solvents under anhydrous conditions, unless otherwise noted. Reagents were purchased at the highest commercial quality and used without any further purification, unless otherwise stated. Room temperature is defined as between 21°C and 25°C. reactions were stirred magnetically and monitored by thin layer chromatography (TLC) using TLC plates precoated with silica gel 60 F254 on aluminum (Merck KGaA). Detection was by UV (254 nm and 365 nm) or chemical stain ($KMnO_4$, ninhydrin, iodine). Solvents were removed *in vacuo* using a Buchi R-300 Rotavapor (equipped with an I-300 Pro Interface, B-300 base Heating Bath, Welch 2037B-01 DryFast pump, and VWR AD15R-40-V11B Circulating Bath). Solvents for silica gel chromatography were used as supplied by Sigma-Aldrich. Automated flash chromatography was performed on a Biotage Isolera instrument, equipped with a UV detector. Chromatograms were recorded at 254 nm and 280 nm. High-resolution mass spectra (HRMS) were obtained using Q Exactive™ Plus Hybrid QuadrupoleOrbitrap™ Mass Spectrometer. $^{1}H$ and $^{13}C$ Nuclear Magnetic Resonance (NMR) spectra were recorded on BRUKER AV (600 MHz and 700 MHz), AVB (400 MHz), AVQ (400 MHz) and NEO (500 MHz) spectrometers. Measurements were carried out at ambient temperature. Chemical shifts ($\delta$) are reported in ppm with the residual solvent signal as internal standard (chloroform at 7.26 and 77.2 ppm for $^{1}H$ NMR and $^{13}C$ NMR, respectively, methanol at 3.31 and 49.0, respectively and DMSO at 2.50 and 39.5, respectively). Multiplicity is reported as follows: singlet (s), doublet (d), doublet of doublet (dd) doublet of triplet (dt), triplet (t), triplet of doublet (td), quartet (q), and multiplet (m). Coupling constants ($J$) are reported in Hertz (Hz). $^{13}C$ NMR spectra were recorded with broadband 1 H decoupling.

**General Procedure A (Amide Coupling):**

To a vessel of amine (1 equiv.), carboxylic acids (1 equiv.) and HATU (1.2 equiv.) was added and purged with $N_2$ for 5 min. DMF (0.2 M) was added followed by adding DIPEA (4 equiv.). The reaction was stirred at ambient temperature overnight. The reaction mixture was dried under air, washed with water, and purified by silica gel flash chromatography to yield the final compound.

**General Procedure B (Boc Deprotecting):**

The tert-butyloxycarbonyl (Boc) protected amine (1 equiv.) was dissolved in DCM (0.1 M). Trifluoroacetic acid (32 equiv.) was added. The reaction mixture was stirred at ambient temperature for 2 hours. The volatiles were removed *in vacuo* and the crude residue was used without further purification.

**Synthetic Scheme for ZD-1-49, 50, 52, 53, 55**

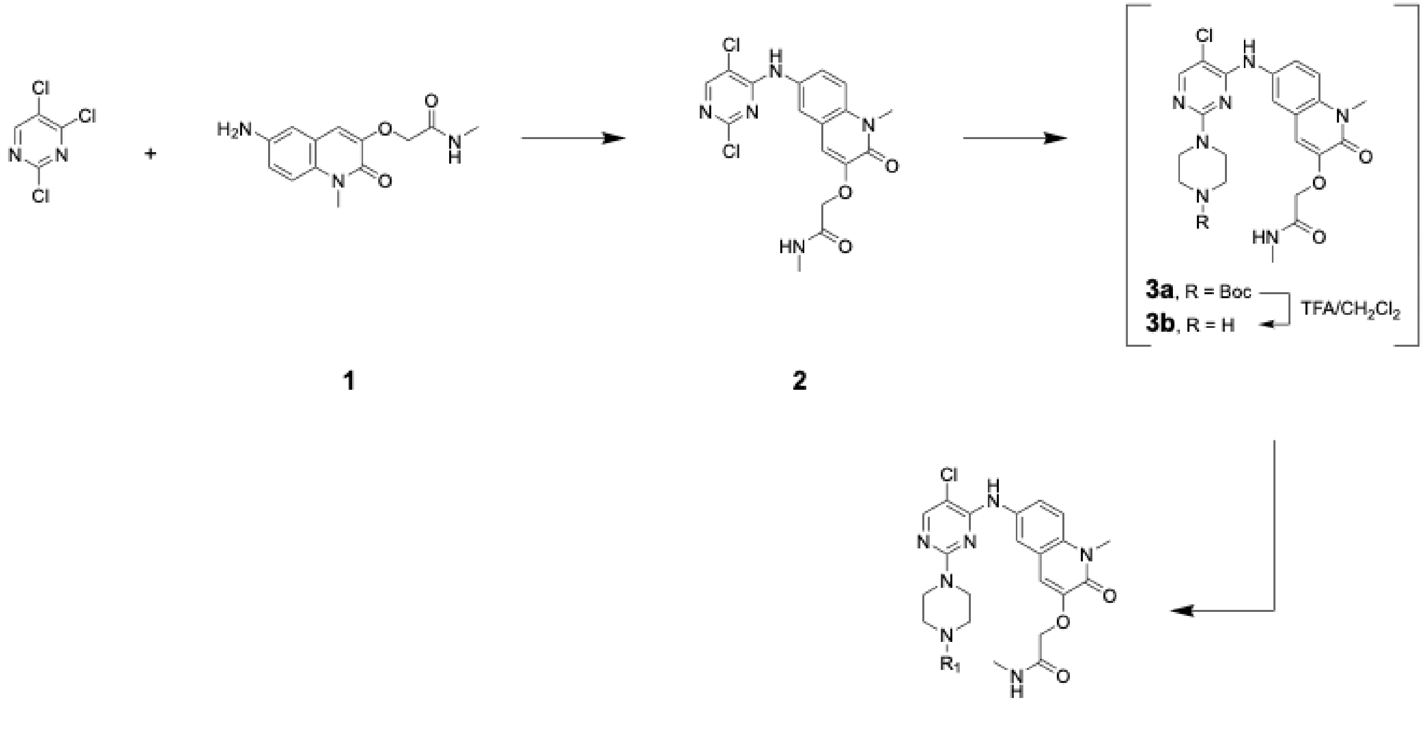


**ZD-1-49, 50, 52, 53, 55**

**Procedure for the preparation of Compound 2[5]:**

To 2-[(6-amino-1-methyl-2-oxo-1,2-dihydroquinolin-3-yl)oxy]-N-methylacetamide (**1**, 2 g, 7.65 mmol) and 2,4,5-trichloropyrimidine (9.19 g, 20.6 mmol) in DMF (10.0 mL) and MeOH (10.0 mL) was added triethylamine (1.86 g, 18.4 mmol) and the mixture was heated to 70 °C overnight. After cooling to room temperature, the reaction mixture was concentrated *in vacuo* and the residue taken up in water. The resulting precipitate was collected by filtration, washed with $Et_2O$ and dried *in vacuo* to give 2-({6-[(2,5-dichloropyrimidin-4-yl)amino]-1-methyl-2-oxo-1,2-dihydroquinolin-3-yl}oxy)-N-methyl-acetamide (**2**, 3.05 g, 7.47 mmol) (70% yield).

***tert*-butyl 4-(5-chloro-4-((1-methyl-3-(2-(methylamino)-2-oxoethoxy)-2-oxo-1,2-dihydroquinolin-6-yl)amino)pyrimidin-2-yl)piperazine-1-carboxylate (3a)**

To 2-({6-[(2,5-dichloropyrimidin-4-yl)amino]-1-methyl-2-oxo-1,2-dihydroquinolin-3-yl}oxy)-N-methyl-acetamide (**2**, 500 mg, 1.23 mmol) and 1-Boc-piperazine (228 mg, 1.23 mmol) in DMF (3 mL) was added triethylamine (496 mg, 4.90 mmol) and the mixture stirred at 100 °C for 2 h. After cooling to room temperature, the mixture was filtered, the reaction mixture was dried under air and purified through silica gel chromatography (0-10% MeOH in DCM) to give *tert*-butyl 4-(5-chloro-4-((1-methyl-3-(2-(methylamino)-2-oxoethoxy)-2-oxo-1,2-dihydroquinolin-6-yl)amino)pyrimidin-2-yl)piperazine-1-carboxylate (**3a**, 611 mg, 79% yield).

$^1$H NMR (500 MHz, DMSO) δ 8.82 (s, 1H), 8.00 (s, 1H), 7.89 (q, *J* = 4.7 Hz, 1H), 7.80 (d, *J* = 2.4 Hz, 1H), 7.69 (dd, *J* = 9.0, 2.5 Hz, 1H), 7.41 (d, *J* = 9.2 Hz, 1H), 7.10 (s, 1H), 4.51 (s, 2H), 3.61 (s, 3H), 3.54 (dd, *J* = 6.9, 3.9 Hz, 4H), 3.31 (dd, *J* = 6.7, 4.1 Hz, 4H), 2.61 (d, *J* = 4.6 Hz, 3H), 1.34 (s, 8H); $^{13}$C NMR (126 MHz, DMSO) δ 167.96, 159.59, 157.17, 155.88, 155.01, 154.39, 147.16, 133.96, 132.31, 123.44, 120.55, 119.99, 114.83, 113.92, 103.00, 79.51, 68.47, 43.98, 30.18, 28.97, 25.89; MS (ESI): Calcd for $C_{26}H_{32}ClN_7O_5^+$ $[M+1]^+$: 557.22; found: 558.23.

**2-((6-((5-chloro-2-(piperazin-1-yl)pyrimidin-4-yl)amino)-1-methyl-2-oxo-1,2-dihydroquinolin-3-yl)oxy)-*N*-methylacetamide (3b)**

Following general procedure B, 2-((6-((5-chloro-2-(piperazin-1-yl)pyrimidin-4-yl)amino)-1-methyl-2-oxo-1,2-dihydroquinolin-3-yl)oxy)-*N*-methylacetamide (**3b**) was obtained from *tert*-butyl 4-(5-chloro-4-((1-methyl-3-(2-(methylamino)-2-oxoethoxy)-2-oxo-1,2-dihydroquinolin-6-yl)amino)pyrimidin-2-yl)piperazine-1-carboxylate (**3a**, 100 mg, 0.18 mmol) as a yellow solid. The intermediate was used without further purification.

**2-((6-((2-(4-acryloylpiperazin-1-yl)-5-chloropyrimidin-4-yl)amino)-1-methyl-2-oxo-1,2-dihydroquinolin-3-yl)oxy)-*N*-methylacetamide (ZD-1-49)**

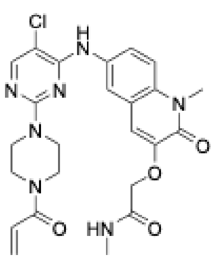


Following general procedure A, 2-((6-((2-(4-acryloylpiperazin-1-yl)-5-chloropyrimidin-4-yl)amino)-1-methyl-2-oxo-1,2-dihydroquinolin-3-yl)oxy)-*N*-methylacetamide (**ZD-1-49**) was obtained from 2-((6-((5-chloro-2-(piperazin-1-yl)pyrimidin-4-yl)amino)-1-methyl-2-oxo-1,2-dihydroquinolin-3-yl)oxy)-*N*-methylacetamide (**3b**, 30 mg, 65.5 μmol) and acryloyl chloride (5.93 mg, 65.5 μmol). The crude residue was purified by silica gel chromatography (0-7% MeOH in DCM) to give 10.5 mg (31% yield) as a yellow-white powder.

$^1$H NMR (500 MHz, DMSO) δ 8.93 (s, 1H), 8.09 (s, 1H), 8.02 (q, *J* = 4.8 Hz, 1H), 7.91 (d, *J* = 2.4 Hz, 1H), 7.76 (dd, *J* = 9.1, 2.5 Hz, 1H), 7.50 (d, *J* = 9.1 Hz, 1H), 7.18 (s, 1H), 6.86 (dd, *J* = 16.7, 10.4 Hz, 1H), 6.15 (dd, *J* = 16.7, 2.4 Hz, 1H), 5.71 (dd, *J* = 10.4, 2.3 Hz, 1H), 4.59 (s, 2H), 3.69 (s, 3H), 3.66 (s, 4H), 3.62 – 3.56 (m, 2H), 2.67 (d, *J* = 4.7 Hz, 3H); $^{13}$C NMR (126 MHz, DMSO) δ 168.05, 164.86, 159.52, 157.20, 155.92, 155.06, 147.11, 133.98, 132.29, 128.63, 128.05, 123.51, 120.63, 120.01, 114.85, 113.89, 103.07, 68.46, 30.21, 25.90; MS (ESI): Calcd for $C_{24}H_{26}ClN_7O_4^+$ [M+1]$^+$:511.17; found: 512.06.

**(*E*)-2-((6-((5-chloro-2-(4-(2-cyano-3-cyclopropylacryloyl)piperazin-1-yl)pyrimidin-4-yl)amino)-1-methyl-2-oxo-1,2-dihydroquinolin-3-yl)oxy)-*N*-methylacetamide (ZD-1-50)**

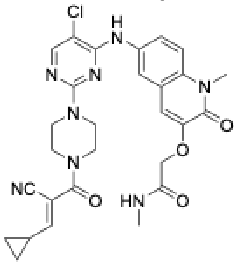


Following general procedure A, (*E*)-2-((6-((5-chloro-2-(4-(2-cyano-3-cyclopropylacryloyl)piperazin-1-yl)pyrimidin-4-yl)amino)-1-methyl-2-oxo-1,2-dihydroquinolin-3-yl)oxy)-*N*-methylacetamide (**ZD-1-50**) was obtained from 2-((6-((5-chloro-2-(piperazin-1-yl)pyrimidin-4-yl)amino)-1-methyl-2-oxo-1,2-dihydroquinolin-3-yl)oxy)-*N*-methylacetamide (**3b**, 21.2 mg, 46.3 μmol) and (*E*)-2-cyano-3-cyclopropylacrylic acid (6.35 mg, 46.3 μmol). The crude residue was purified by silica gel chromatography (0-7% MeOH in DCM) to give 12 mg (45% yield) as a yellow-white powder.

$^1$H NMR (500 MHz, DMSO) δ 8.92 (d, *J* = 6.7 Hz, 1H), 8.09 (d, *J* = 3.7 Hz, 1H), 7.98 (t, *J* = 4.9 Hz, 1H), 7.87 (t, *J* = 2.4 Hz, 1H), 7.78 (dt, *J* = 9.1, 2.6 Hz, 1H), 7.52 – 7.45 (m, 1H), 7.19 (s, 1H), 6.63 (d, *J* = 11.0 Hz, 1H), 6.41 (d, *J* = 11.2 Hz, 0H), 4.58 (s, 2H), 3.69 (d, *J* = 1.2 Hz, 4H), 3.61 – 3.55 (m, 3H), 3.37 (s, 4H), 2.68 (d, *J* = 4.7 Hz, 3H), 1.90 (ddq, *J* = 15.9, 8.1, 4.1 Hz, 1H), 1.20 (tq, *J* = 7.1, 4.1 Hz, 2H), 0.96 – 0.90 (m, 1H), 0.88 – 0.80 (m, 1H); $^{13}$C NMR (126 MHz, DMSO) δ 167.98, 164.42, 162.38, 159.50, 157.19, 155.92, 155.01, 147.12, 133.95, 132.32, 123.49, 120.55, 120.03, 116.22, 114.86, 113.97, 106.65, 103.21, 68.49, 30.19, 25.89, 15.91, 10.49; MS (ESI): Calcd for $C_{28}H_{29}ClN_8O_4^+$ [M+1]$^+$: 576.20; found: 577.20.

**2-((6-((2-(4-(3-bromo-4,5-dihydroisoxazole-5-carbonyl)piperazin-1-yl)-5-chloropyrimidin-4-yl)amino)-1-methyl-2-oxo-1,2-dihydroquinolin-3-yl)oxy)-*N*-methylacetamide (ZD-1-52)**

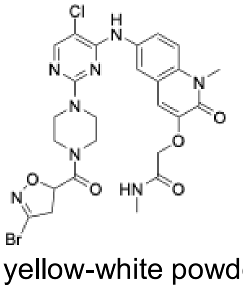


A stirred solution of 2-((6-((5-chloro-2-(piperazin-1-yl)pyrimidin-4-yl)amino)-1-methyl-2-oxo-1,2-dihydroquinolin-3-yl)oxy)-*N*-methylacetamide (**3b**, 30 mg, 65.5 μmol**)** and HATU (1.2 equiv.) in DMF (0.2 M) was purged with $N_2$ for 5 min. 3-bromo-4,5-dihydroisoxazole-5-carboxylic acid (12.71 mg, 65.5 μmol) in DMF and DIPEA (4 equiv.) was added and stirred at ambient temperature overnight. The crude residue was purified by silica gel chromatography (0-7% MeOH in DCM) to give 2-((6-((2-(4-(3-bromo-4,5-dihydroisoxazole-5-carbonyl)piperazin-1-yl)-5-chloropyrimidin-4-yl)amino)-1-methyl-2-oxo-1,2-dihydroquinolin-3-yl)oxy)-*N*-methylacetamide (**ZD-1-52**) (5.4 mg, 13% yield) as a yellow-white powder.

$^1$H NMR (500 MHz, DMSO) δ 8.96 (s, 1H), 8.11 (s, 1H), 8.01 (s, 1H), 7.86 (s, 1H), 7.78 (d, *J* = 9.0 Hz, 1H), 7.50 (d, *J* = 9.0 Hz, 1H), 7.20 (s, 1H), 4.58 (s, 2H), 3.76 – 3.71 (m, 4H), 3.69 (s, 3H), 3.65 (s, 4H), 2.67 (d, *J* = 4.7 Hz, 3H), 1.66 (d, *J* = 35.9 Hz, 1H), 1.50 (d, 1H), 0.86 (s, 1H); $^{13}$C NMR (126 MHz, DMSO) δ 168.02, 165.89, 159.53, 157.18, 155.94, 155.05, 147.15, 138.91, 133.95, 132.35, 123.53, 120.64, 120.01, 114.83, 113.94, 103.16, 77.70, 68.49, 30.19, 25.90; MS (ESI): Calcd for $C_{25}H_{26}BrClN_8O_5^+$ [M+1]$^+$: 632.08; found: 633.12.

**2-((6-((2-(4-(2-(1-acryloylazetidin-3-yl)acetyl)piperazin-1-yl)-5-chloropyrimidin-4-yl)amino)-1-methyl-2-oxo-1,2-dihydroquinolin-3-yl)oxy)-*N*- methylacetamide (ZD-1-53)**

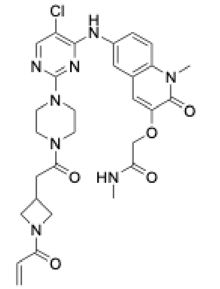

Following general procedure A, 2-((6-((2-(4-(2-(1-acryloylazetidin-3-yl)acetyl)piperazin-1-yl)-5-chloropyrimidin-4-yl)amino)-1-methyl-2-oxo-1,2-dihydroquinolin-3-yl)oxy)-*N*- methylacetamide (**ZD-1-53**) was obtained from 2-((6-((5-chloro-2-(piperazin-1-yl)pyrimidin-4-yl)amino)-1-methyl-2-oxo-1,2-dihydroquinolin-3-yl)oxy)-*N*-methylacetamide (**3b**, 21.2 mg, 46.3 µmol) and 2-(1-acryloylazetidin-3-yl)acetic acid (7.84 mg, 46.3 µmol). The crude residue was purified by silica gel chromatography (0-7% MeOH in DCM) to give 13.4 mg (48% yield) as a yellow-white powder.

$^1$H NMR (500 MHz, DMSO) δ 8.91 (s, 1H), 8.08 (s, 1H), 8.00 (q, *J* = 4.7 Hz, 1H), 7.91 (d, *J* = 2.4 Hz, 1H), 7.76 (dd, *J* = 9.1, 2.4 Hz, 1H), 7.48 (d, *J* = 9.1 Hz, 1H), 7.16 (s, 1H), 6.28 (dd, *J* = 17.0, 10.3 Hz, 1H), 6.07 (dd, *J* = 17.0, 2.3 Hz, 1H), 5.63 (dd, *J* = 10.3, 2.3 Hz, 1H), 4.59 (s, 2H), 4.32 (t, *J* = 8.5 Hz, 1H), 4.01 (dd, *J* = 10.1, 8.3 Hz, 1H), 3.83 (dd, *J* = 8.7, 5.5 Hz, 1H), 3.68 (s, 5H), 3.61 (t, *J* = 5.4 Hz, 2H), 3.56 (dd, *J* = 10.1, 5.7 Hz, 1H), 3.53 – 3.49 (m, 4H), 2.94 – 2.82 (m, 1H), 2.78 (d, *J* = 7.7 Hz, 2H), 2.67 (d, *J* = 4.7 Hz, 3H); $^{13}$C NMR (126 MHz, DMSO) δ 169.82, 168.03, 164.94, 159.55, 157.18, 155.88, 155.04, 147.11, 134.00, 132.25, 127.46, 126.56, 123.42, 120.54, 119.99, 114.80, 113.84, 103.03, 68.46, 55.61, 53.09, 30.19, 25.90, 25.58; MS (ESI): Calcd for $C_{29}H_{33}ClN_8O_5^+$ [M+1]$^+$: 608.22; found: 609.23.

**2-((6-((2-(4-(bicyclo[1.1.0]butane-1-carbonyl)piperazin-1-yl)-5-chloropyrimidin-4-yl)amino)-1-methyl-2-oxo-1,2-dihydroquinolin-3-yl)oxy)-*N*-methylacetamide (ZD-1-55)**

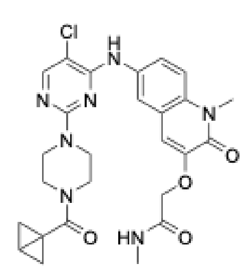

Following general procedure A, 2-((6-((2-(4-(bicyclo[1.1.0]butane-1-carbonyl)piperazin-1-yl)-5-chloropyrimidin-4-yl)amino)-1-methyl-2-oxo-1,2-dihydroquinolin-3-yl)oxy)-*N*-methylacetamide (**ZD-1-55**) was obtained from 2-((6-((5-chloro-2-(piperazin-1-yl)pyrimidin-4-yl)amino)-1-methyl-2-oxo-1,2-dihydroquinolin-3-yl)oxy)-*N*-methylacetamide (**3b**, 30 mg, 65.5 µmol) and sodium bicyclo[1.1.0]butane-1-carboxylate (6.36 mg, 65.5 µmol). The crude residue was purified by silica gel chromatography (0-7% MeOH in DCM) to give 12.6 mg (36% yield) as a yellow-white powder.

$^1$H NMR (500 MHz, DMSO) δ 8.91 (t, *J* = 1.6 Hz, 1H), 8.10 (dt, *J* = 8.6, 1.0 Hz, 1H), 7.98 (q, *J* = 4.6 Hz, 1H), 7.94 – 7.84 (m, 1H), 7.78 (dt, *J* = 9.0, 2.4 Hz, 1H), 7.50 (dd, *J* = 9.0, 1.5 Hz, 1H), 7.19 (d, *J* = 6.5 Hz, 1H), 4.58 (s, 2H), 3.74 (dd, *J* = 6.8, 3.7 Hz, 1H), 3.69 (d, *J* = 1.8 Hz, 3H), 3.66 (d, *J* = 5.1 Hz, 3H), 3.61 (s, 1H), 3.35 (s, 5H), 2.67 (d, *J* = 4.6 Hz, 3H), 2.20 (d, *J* = 3.3 Hz, 1H), 1.99 (p, *J* = 3.0 Hz, 1H), 1.24 (d, *J* = 11.7 Hz, 1H), 1.13 (d, *J* = 2.5 Hz, 1H); $^{13}$C NMR (126 MHz, DMSO) δ 169.70, 167.96, 159.57, 157.20, 155.91, 155.04, 155.01, 147.10, 133.99, 132.29, 123.48, 120.57, 120.03, 114.90, 113.93, 103.05, 68.45, 37.30, 30.21, 25.85, 13.54, 8.21; MS (ESI): Calcd for $C_{26}H_{28}ClN_7O_4^+$ [M+1]$^+$: 537.18; found 538.15.

**Synthetic Scheme for ZD-1-181, 182, 184, 186**

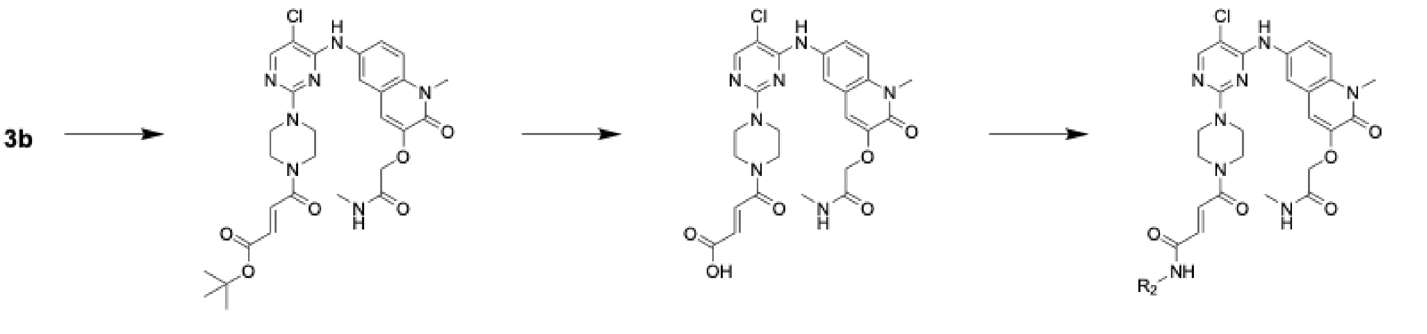

**4** **5** **ZD-1-181, 182, 184, 186**

***tert*-butyl (*E*)-4-(4-(5-chloro-4-((1-methyl-3-(2-(methylamino)-2-oxoethoxy)-2-oxo-1,2-dihydroquinolin-6-yl)amino)pyrimidin-2-yl)piperazin-1-yl)-4-oxobut-2-enoate (4)**

Following general procedure A, *tert*-butyl (*E*)-4-(4-(5-chloro-4-((1-methyl-3-(2-(methylamino)-2-oxoethoxy)-2-oxo-1,2-dihydroquinolin-6-yl)amino)pyrimidin-2-yl)piperazin-1-yl)-4-oxobut-2-enoate (**4**) was obtained from 2-((6-((5-chloro-2-(piperazin-1-yl)pyrimidin-4-yl)amino)-1-methyl-2-oxo-1,2-dihydroquinolin-3-yl)oxy)-*N*-methylacetamide (**3b**, 84.8 mg, 0.185 mmol) and (*E*)-4-(*tert*-butoxy)-4-oxobut-2-enoic acid (31.9 mg, 0.185 mmol). The crude residue was purified by silica gel chromatography (0-7% MeOH in DCM) to give 75.6 mg (67% yield) as a yellow-white powder.

$^1$H NMR (500 MHz, DMSO) δ 8.92 (s, 1H), 8.09 (d, *J* = 1.2 Hz, 1H), 7.88 (d, *J* = 2.4 Hz, 1H), 7.76 (dd, *J* = 9.1, 2.4 Hz, 1H), 7.49 (d, *J* = 9.1 Hz, 1H), 7.40 (d, *J* = 15.4 Hz, 1H), 7.18 (s, 1H), 6.49 (d, *J* = 15.3 Hz, 1H), 4.58 (s, 1H), 3.68 (s, 3H), 3.66 (s, 3H), 3.65 – 3.59 (m, 2H), 3.42 (s, 5H), 2.66 (s, 1H), 1.46 (s, 9H); MS (ESI): Calcd for $C_{29}H_{34}ClN_7O_6^+$ [M+1]$^+$: 611.22; found 612.54.

**(*E*)-4-(4-(5-chloro-4-((1-methyl-3-(2-(methylamino)-2-oxoethoxy)-2-oxo-1,2-dihydroquinolin-6-yl)amino)pyrimidin-2-yl)piperazin-1-yl)-4-oxobut-2-enoic acid (5)**

Following general procedure B, (*E*)-4-(4-(5-chloro-4-((1-methyl-3-(2-(methylamino)-2-oxoethoxy)-2-oxo-1,2-dihydroquinolin-6-yl)amino)pyrimidin-2-yl)piperazin-1-yl)-4-oxobut-2-enoic acid (**5**) was obtained from **4** (50 mg, 0.089 mmol) as a yellow solid. The intermediate was used without further purification.

**(*E*)-4-(4-(5-chloro-4-((1-methyl-3-(2-(methylamino)-2-oxoethoxy)-2-oxo-1,2-dihydroquinolin-6-yl)amino)pyrimidin-2-yl)piperazin-1-yl)-*N*-((3-(4-fluorophenyl)-1,2,4-oxadiazol-5-yl)methyl)-4-oxobut-2-enamide (ZD-1-181)**

Following general procedure A, (*E*)-4-(4-(5-chloro-4-((1-methyl-3-(2-(methylamino)-2-oxoethoxy)-2-oxo-1,2-dihydroquinolin-6-yl)amino)pyrimidin-2-yl)piperazin-1-yl)-*N*-((3-(4-fluorophenyl)-1,2,4-oxadiazol-5-yl)methyl)-4-oxobut-2-enamide (**ZD-1-181**) was obtained from **5** (20.5 mg, 36.8 μmol) and (3-(4-fluorophenyl)-1,2,4-oxadiazol-5-yl)methanamine (7.11 mg, 36.8 μmol). The crude residue was purified by silica gel chromatography (0-7% MeOH in DCM) to give 10.7 mg (40% yield) as a yellow-white powder.

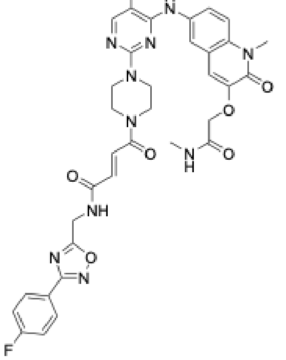

$^1$H NMR (500 MHz, DMSO) δ 9.42 (q, *J* = 7.1 Hz, 1H), 8.18 (d, *J* = 2.1 Hz, 1H), 8.08 – 7.98 (m, 3H), 7.84 (t, *J* = 2.7 Hz, 1H), 7.75 (dt, *J* = 9.0, 2.8 Hz, 1H), 7.52 (d, *J* = 9.0 Hz, 1H), 7.45 – 7.36 (m, 3H), 7.21 (s, 1H), 6.93 (d, *J* = 15.1 Hz, 1H), 4.77 (d, *J* = 5.6 Hz, 2H), 4.59 (s, 2H), 3.70 (s, 7H), 3.69 (s, 4H), 3.67 – 3.62 (m, 4H), 2.66 (d, *J* = 4.6 Hz, 3H); $^{13}$C NMR (126 MHz, DMSO) δ 178.01, 167.95, 167.33, 165.44, 164.89, 163.80, 163.46, 157.22, 156.46, 147.21, 133.99, 132.93, 130.70, 130.01, 123.05, 120.08, 116.88, 114.96, 113.74, 68.41, 45.03, 44.35, 41.54, 35.89, 30.25, 25.87; MS (ESI): Calcd for $C_{34}H_{32}ClFN_{10}O_6^+$ [M+1]$^+$: 730.21; found 731.25.

**(*E*)-4-(4-(5-chloro-4-((1-methyl-3-(2-(methylamino)-2-oxoethoxy)-2-oxo-1,2-dihydroquinolin-6-yl)amino)pyrimidin-2-yl)piperazin-1-yl)-*N*-((1,3-dimethyl-2-oxo-2,3-dihydro-1*H*-benzo[*d*]imidazol-5-yl)methyl)-4-oxobut-2-enamide (ZD-1-182)**

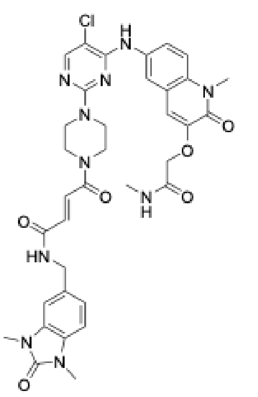

Following general procedure A, (*E*)-4-(4-(5-chloro-4-((1-methyl-3-(2-(methylamino)-2-oxoethoxy)-2-oxo-1,2-dihydroquinolin-6-yl)amino)pyrimidin-2-yl)piperazin-1-yl)-*N*-((1,3-dimethyl-2-oxo-2,3-dihydro-1*H*-benzo[*d*]imidazol-5-yl)methyl)-4-oxobut-2-enamide (**ZD-1-182**) was obtained from **5** (18.6 mg, 33.4 μmol) and 5-(aminomethyl)-1,3-dimethyl-1,3-dihydro-2*H*-benzo[*d*]imidazol-2-one (6.39 mg, 33.4 μmol). The crude residue was purified by silica gel chromatography (0-7% MeOH in DCM) to give 12 mg (49% yield) as a yellow-white powder.

$^1$H NMR (500 MHz, DMSO) δ 8.94 – 8.86 (m, 2H), 8.09 (d, *J* = 1.1 Hz, 1H), 7.98 (q, *J* = 4.3 Hz, 1H), 7.88 (d, *J* = 2.4 Hz, 1H), 7.79 (dd, *J* = 9.1, 2.4 Hz, 1H), 7.50 (d, *J* = 9.1 Hz, 1H), 7.35 (d, *J* = 15.1 Hz, 1H), 7.19 (s, 1H), 7.12 – 7.05 (m, 2H), 7.00 (dd, *J* = 8.1, 1.6 Hz, 1H), 6.91 (d, *J* = 15.0 Hz, 1H), 4.59 (s, 2H), 4.42 (d, *J* = 5.8 Hz, 2H), 3.69 (s, 6H), 3.67 (s, 4H), 3.64 – 3.60 (m, 2H), 3.35 (s, 3H), 3.31 (d, *J* = 1.5 Hz, 3H), 2.67 (d, *J* = 4.7 Hz, 3H); $^{13}$C NMR (126 MHz, DMSO) δ 168.02, 163.97, 159.50, 157.19, 155.92, 155.05, 154.45, 147.15, 135.31, 133.97, 132.24, 130.20, 129.56, 129.27, 123.52, 120.75, 120.55, 120.02, 114.84, 114.01, 107.83, 107.41, 103.09, 68.49, 43.16, 30.19, 27.38, 25.89; MS (ESI): Calcd for $C_{35}H_{37}ClN_{10}O_6^+$ [M+1]$^+$: 728.25; found 729.26.

**(*E*)-4-(4-(5-chloro-4-((1-methyl-3-(2-(methylamino)-2-oxoethoxy)-2-oxo-1,2-dihydroquinolin-6-yl)amino)pyrimidin-2-yl)piperazin-1-yl)-*N*-(5-(3-methoxyphenyl)-1,3,4-thiadiazol-2-yl)-4-oxobut-2-enamide (ZD-1-184)**

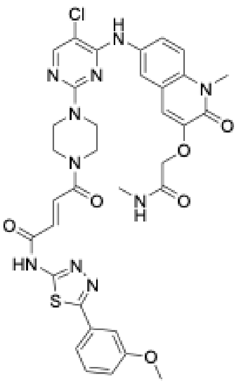

Following general procedure A, (*E*)-4-(4-(5-chloro-4-((1-methyl-3-(2-(methylamino)-2-oxoethoxy)-2-oxo-1,2-dihydroquinolin-6-yl)amino)pyrimidin-2-yl)piperazin-1-yl)-*N*-(5-(3-methoxyphenyl)-1,3,4-thiadiazol-2-yl)-4-oxobut-2-enamide (**ZD-1-184**) was obtained from **5** (12.4 mg, 22.2 µmol) and 5-(3-methoxyphenyl)-1,3,4-thiadiazol-2-amine (4.51 mg, 22.2 µmol). The crude residue was purified by silica gel chromatography (0-7% MeOH in DCM) to give 8.2 mg (49% yield) as a yellow-white powder.

$^1$H NMR (500 MHz, DMSO) δ 13.16 (s, 1H), 8.98 (d, *J* = 5.9 Hz, 1H), 8.10 (d, *J* = 4.7 Hz, 1H), 8.00 (q, *J* = 4.8 Hz, 1H), 7.88 (dd, *J* = 6.6, 2.5 Hz, 1H), 7.78 (td, *J* = 8.3, 2.5 Hz, 1H), 7.67 (d, *J* = 15.2 Hz, 1H), 7.54 – 7.43 (m, 3H), 7.22 – 7.16 (m, 1H), 7.15 – 7.08 (m, 1H), 4.59 (d, *J* = 3.4 Hz, 2H), 3.86 (s, 2H), 3.73 – 3.61 (m, 10H), 2.67 (d, *J* = 4.7 Hz, 3H); $^{13}$C NMR (125 MHz, DMSO-d6): δ 167.9, 157.9, 157.4, 157.2, 155.8, 147.2, 133.5, 132.5, 124.8, 122.4, 120.2, 115.2, 114.0, 113.6, 68.4, 30.2, 25.9; MS (ESI): Calcd for $C_{34}H_{33}ClN_{10}O_6S^+$ [M+1]$^+$: 744.19; found 745.20.

**(*E*)-5-(4-(5-chloro-4-((1-methyl-3-(2-(methylamino)-2-oxoethoxy)-2-oxo-1,2-dihydroquinolin-6-yl)amino)pyrimidin-2-yl)piperazin-1-yl)-*N*-(3-fluoro-4-thiomorpholinophenyl)-4-oxopent-2-enamide (ZD-1-186)**

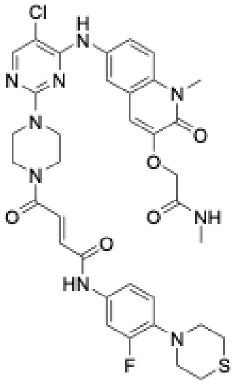

Following general procedure A, (*E*)-5-(4-(5-chloro-4-((1-methyl-3-(2-(methylamino)-2-oxoethoxy)-2-oxo-1,2-dihydroquinolin-6-yl)amino)pyrimidin-2-yl)piperazin-1-yl)-*N*-(3-fluoro-4-thiomorpholinophenyl)-4-oxopent-2-enamide (**ZD-1-186**) was obtained from **5** (92.8 mg, 0.106 mmol) and 3-fluoro-4-thiomorpholinoaniline (22.5 mg, 0.106 µmol). The crude residue was purified by silica gel chromatography (0-7% MeOH in DCM) to give 62.5 mg (78% yield) as a yellow-white powder.

$^1$H NMR (500 MHz, DMSO) δ 10.50 (s, 1H), 8.92 (s, 1H), 8.09 (s, 1H), 7.97 (q, *J* = 4.7 Hz, 1H), 7.88 (d, *J* = 2.5 Hz, 1H), 7.79 (dd, *J* = 9.1, 2.4 Hz, 1H), 7.63 (dd, *J* = 14.4, 2.4 Hz, 1H), 7.50 (d, *J* = 9.2 Hz, 1H), 7.46 (d, *J* = 14.9 Hz, 1H), 7.33 – 7.27 (m, 1H), 7.19 (s, 1H), 7.10 – 7.02 (m, 2H), 4.60 (s, 2H), 3.71 (s, 5H), 3.69 (s, 3H), 3.68 (s, 1H), 3.67 – 3.62 (m, 2H), 3.23 – 3.17 (m, 4H), 2.77 – 2.71 (m, 4H), 2.68 (d, *J* = 4.7 Hz, 3H); $^{19}$F NMR (471 MHz, DMSO) δ -69.29, -70.81; $^{13}$C NMR (126 MHz, DMSO) δ 168.02, 163.69, 162.49, 159.50, 157.19, 155.93, 155.05, 154.04, 147.16, 137.20, 137.13, 135.37, 134.61, 134.53, 133.97, 132.35, 130.50, 123.53, 121.11, 120.57, 120.02, 115.89, 114.84, 114.04, 108.11, 107.90, 103.10, 68.52, 53.52, 30.19, 27.69, 25.89; MS (ESI): Calcd for $C_{35}H_{37}ClFN_9O_5S^+$ [M+1]$^+$: 749.23; found 750.23.

**Synthetic Scheme of Non-reactive version of ZD-1-186**

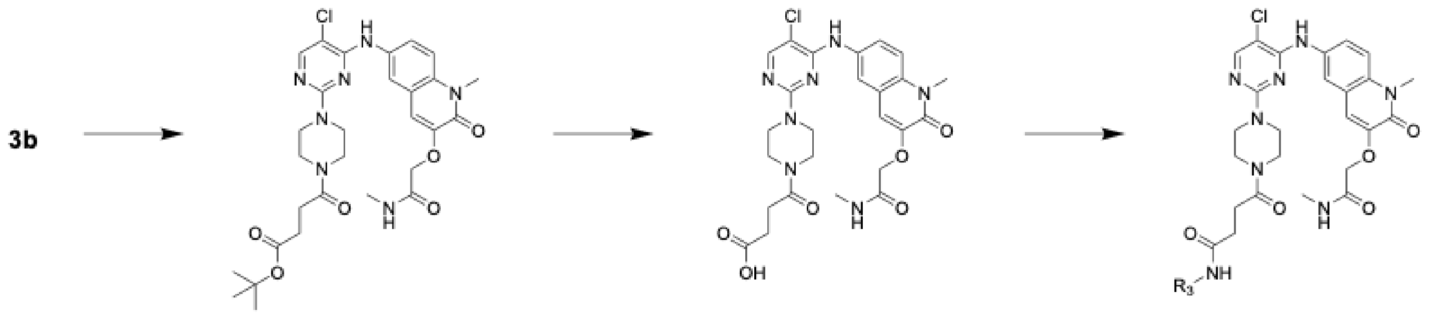

6 7 ZD-2-2

**tert-butyl 4-(4-(5-chloro-4-((1-methyl-3-(2-(methylamino)-2-oxoethoxy)-2-oxo-1,2-dihydroquinolin-6-yl)amino)pyrimidin-2-yl)piperazin-1-yl)-4-oxobutanoate (6)**

Following general procedure A, *tert*-butyl 4-(4-(5-chloro-4-((1-methyl-3-(2-(methylamino)-2-oxoethoxy)-2-oxo-1,2-dihydroquinolin-6-yl)amino)pyrimidin-2-yl)piperazin-1-yl)-4-oxobutanoate (**6**) was obtained from 2-((6-((5-chloro-2-(piperazin-1-yl)pyrimidin-4-yl)amino)-1-methyl-2-oxo-1,2-dihydroquinolin-3-yl)oxy)-*N*-methylacetamide (**3b**, 85.5 mg, 0.187 mmol) and 4-(*tert*-butoxy)-4-oxobutanoic acid (32.5 mg, 0.186 mmol). The crude residue was purified by silica gel chromatography (0-7% MeOH in DCM) to give 70 mg (61% yield) as a yellow-white powder.

$^1$H NMR (400 MHz, DMSO-$D_6$) δ 8.87 (s, 1H), 8.04 (s, 1H), 7.92 (q, *J* = 4.7 Hz, 1H), 7.86 (d, *J* = 2.4 Hz, 1H), 7.73 (dd, *J* = 9.1, 2.4 Hz, 1H), 7.45 (d, *J* = 9.1 Hz, 1H), 7.13 (s, 1H), 4.54 (s, 2H), 3.64 (s, 5H), 3.56 (d, *J* = 5.5 Hz, 2H), 3.49 (q, *J* = 5.1 Hz, 4H), 2.63 (d, *J* = 4.6 Hz, 3H), 2.53 (dd, *J* = 7.5, 5.4 Hz, 2H), 2.38 (dd, *J* = 7.3, 5.4 Hz, 2H), 1.34 (s, 9H); MS (ESI): Calcd for $C_{29}H_{36}ClN_7O_6^+$ [M+1]$^+$: 613.24; found 614.24.

**4-(4-(5-chloro-4-((1-methyl-3-(2-(methylamino)-2-oxoethoxy)-2-oxo-1,2-dihydroquinolin-6-yl)amino)pyrimidin-2-yl)piperazin-1-yl)-4-oxobutanoic acid (7)**

Following general procedure B, 4-(4-(5-chloro-4-((1-methyl-3-(2-(methylamino)-2-oxoethoxy)-2-oxo-1,2-dihydroquinolin-6-yl)amino)pyrimidin-2-yl)piperazin-1-yl)-4-oxobutanoic acid (**7**) was obtained from **6** (63.2 mg, 0.103 mmol) as a yellow solid. The intermediate was used without further purification.

**5-(4-(5-chloro-4-((1-methyl-3-(2-(methylamino)-2-oxoethoxy)-2-oxo-1,2-dihydroquinolin-6-yl)amino)pyrimidin-2-yl)piperazin-1-yl)-*N*-(3-fluoro-4-thiomorpholinophenyl)-4-oxopentanamide (ZD-2-2)**

Following general procedure A, 5-(4-(5-chloro-4-((1-methyl-3-(2-(methylamino)-2-oxoethoxy)-2-oxo-1,2-dihydroquinolin-6-yl)amino)pyrimidin-2-yl)piperazin-1-yl)-*N*-(3-fluoro-4-thiomorpholinophenyl)-4-oxopentanamide (**ZD-2-2**) was obtained from **7** (57.42 mg, 0.103 mmol) and 3-fluoro-4-thiomorpholinoaniline (21.8 mg, 0.103 mmol). The crude residue was purified by silica gel chromatography (0-7% MeOH in DCM) to give 46 mg (59% yield) as a yellow-white powder.

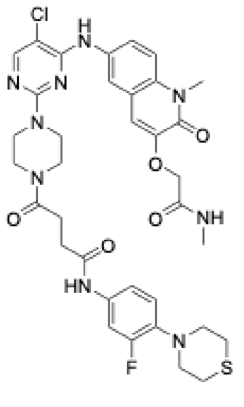


$^1$H NMR (500 MHz, DMSO-d6): δ 9.67 (s, 1H), 8.39 (s, 1H), 7.95 (brq, J = 4.4 Hz, 1H), 7.75 (d, J = 2.2 Hz, 1H), 7.67 (dd, J = 2.2, 9.1 Hz, 1H), 7.54 (d, J = 9.1 Hz, 1H), 7.24 (s, 1H), 4.59 (s, 2H), 3.70 (s, 3H), 2.67 (d, J = 4.4 Hz, 3H); $^{13}$C NMR (126 MHz, DMSO) δ 170.96, 170.47, 168.02, 159.57, 157.18, 155.89, 155.87, 155.03, 154.17, 147.13, 136.22, 135.51, 134.00, 132.29, 123.46, 121.00, 120.98, 120.53, 120.01, 115.22, 114.80, 113.95, 103.03, 68.48, 53.66, 30.18, 27.76, 25.89; MS (ESI): Calcd for $C_{35}H_{39}ClFN_9O_5S^+$ [M+1]$^+$: 751.24; found 752.25.

**3a**

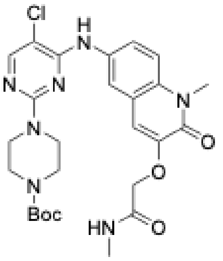

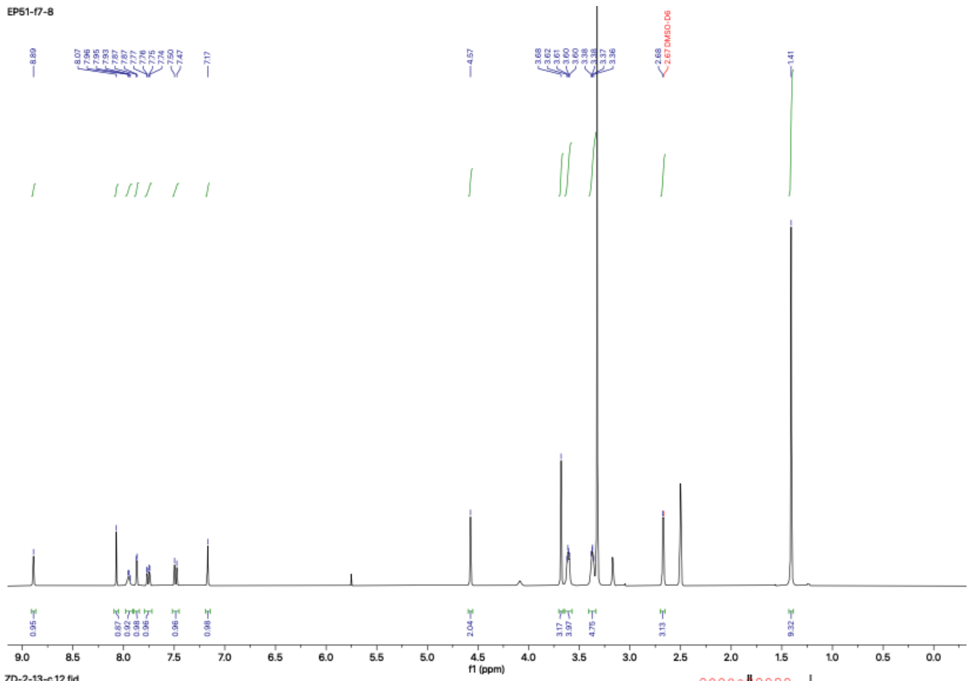


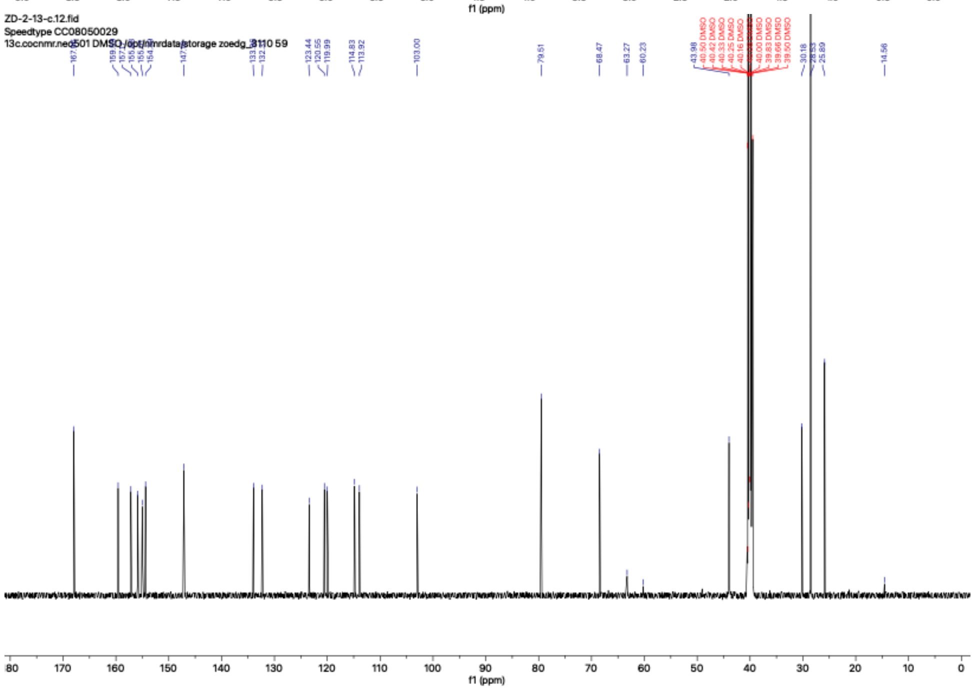

**ZD-1-49**

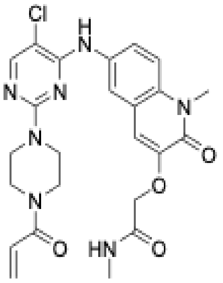

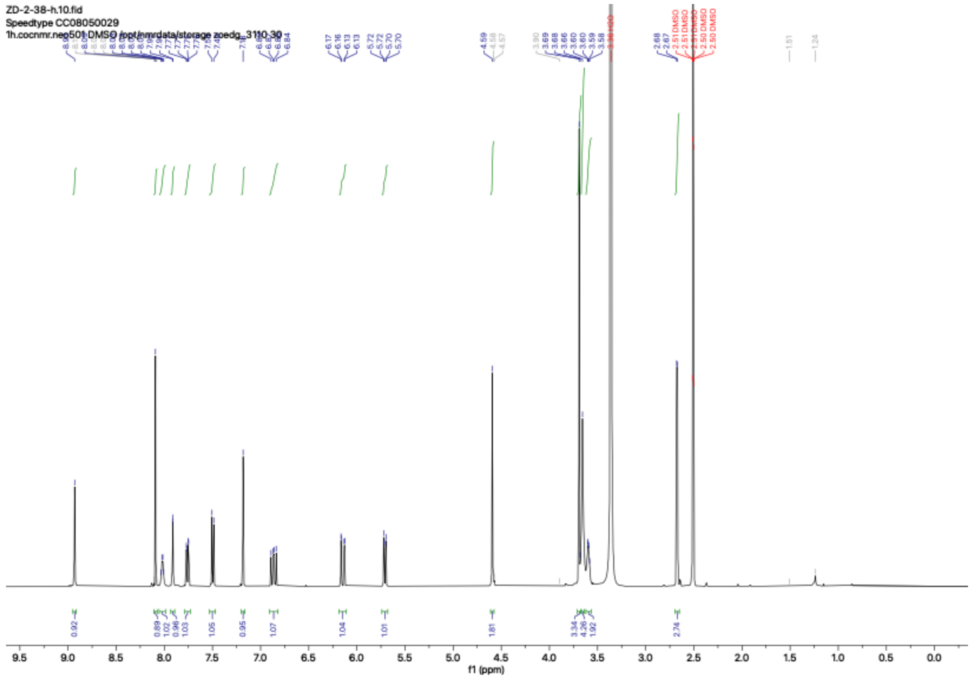

ZD-2-38-h.10.fid
Speedtype CC08050029
1h.cocnmr.neo501 DMSO /opt/nmrdata/storage zoedg_3110 30
f1 (ppm)


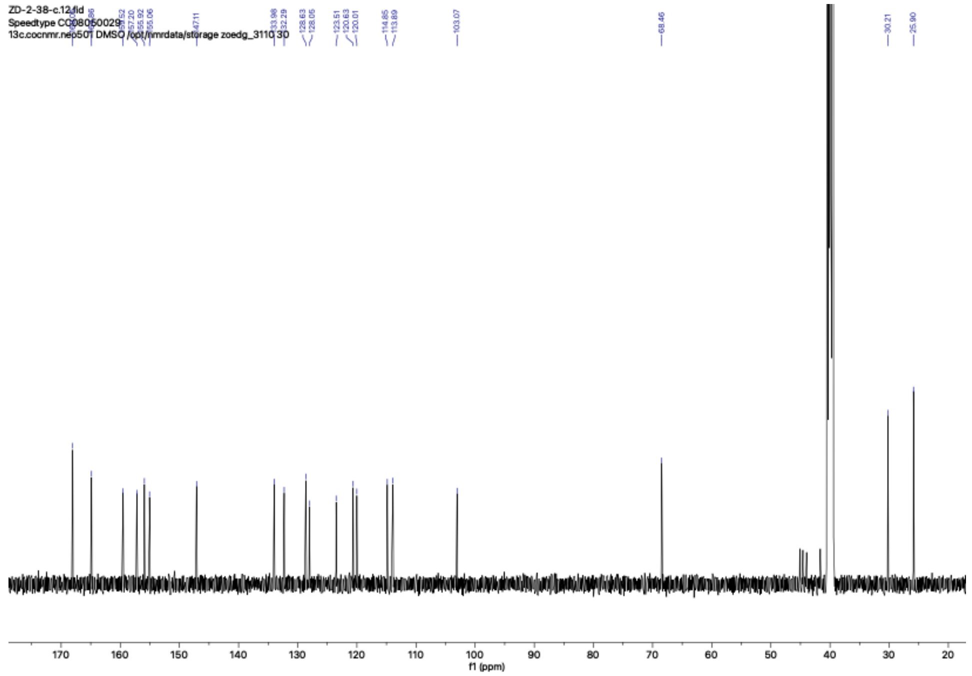

ZD-2-38-c.12.fid
Speedtype CC08050029
13c.cocnmr.neo501 DMSO /opt/nmrdata/storage zoedg_3110 30
133.98
132.29
128.63
128.05
123.51
120.63
120.01
114.85
113.89
103.07
68.46
30.21
25.90
f1 (ppm)

**ZD-1-50**

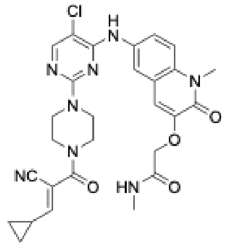
Cl
N
H
N
N
N
N
N
O
NC
O
O
HN
O

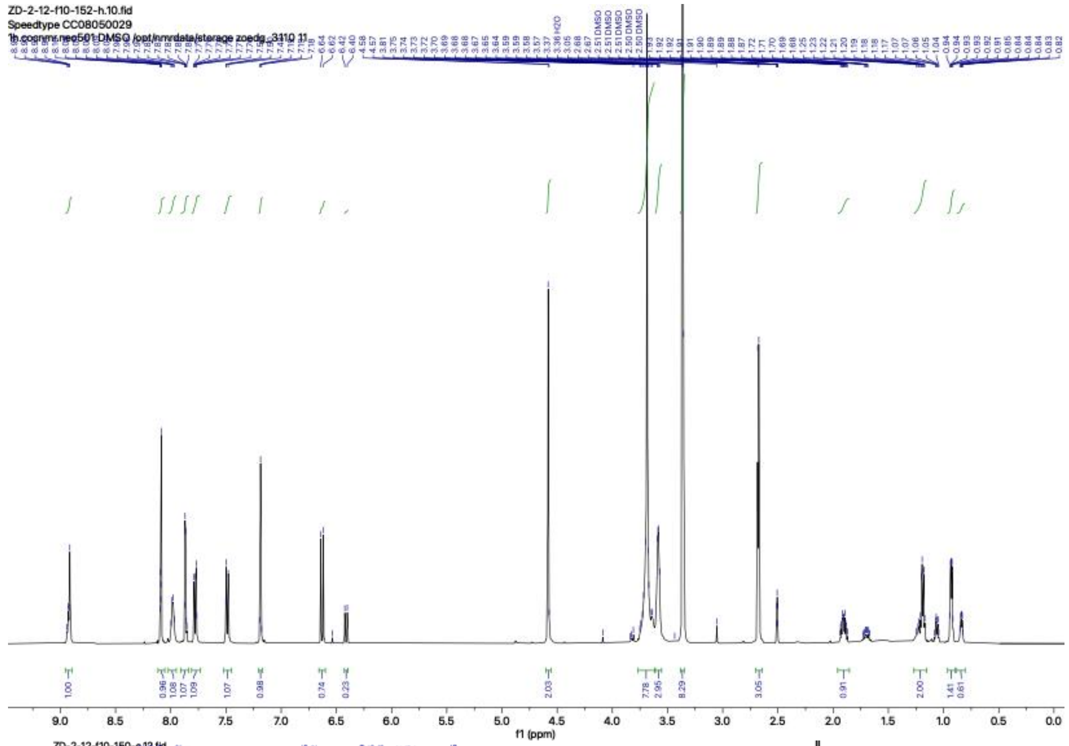
ZD-2-12-f10-152-h.10.fid
Speedtype CC08050029
1h.cocnmr.neo501 DMSO /opt/nmrdata/storage zoedg_3110 11
3.37 H2O
2.51 DMSO
2.50 DMSO
f1 (ppm)

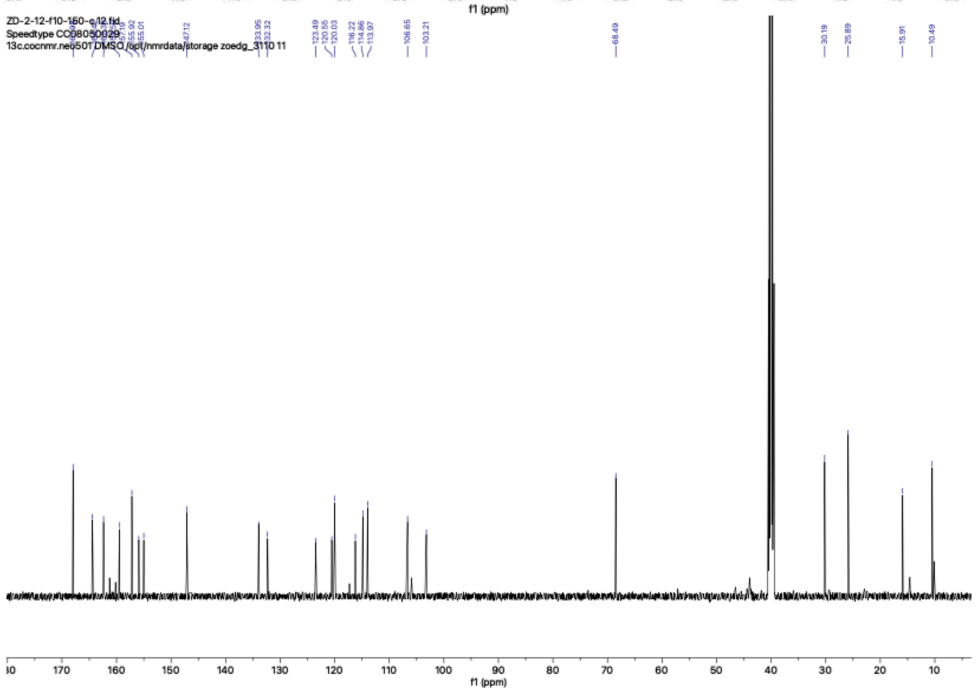
ZD-2-12-f10-150-c.12.fid
Speedtype CC08050029
13c.cocnmr.neo501 DMSO /opt/nmrdata/storage zoedg_3110 11
123.49
120.55
120.03
116.22
114.86
113.97
106.65
103.21
68.49
30.19
25.89
15.91
10.49
f1 (ppm)

**ZD-1-52**

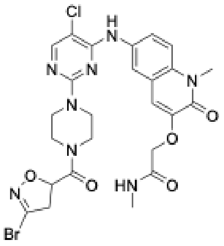

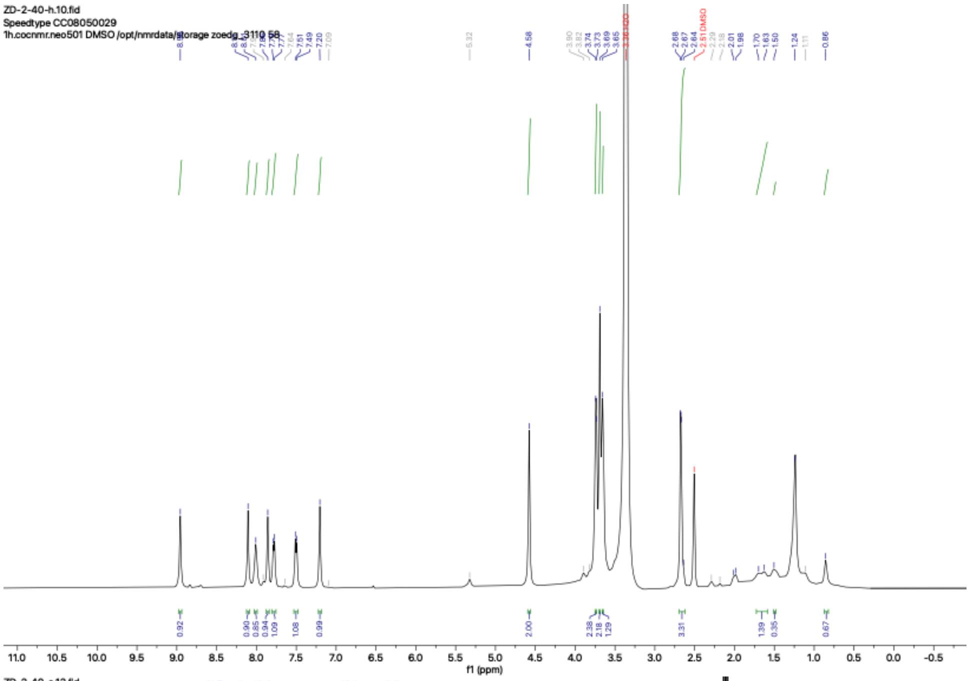

ZD-2-40-h.10.fid
Speedtype CC08050029
1h.cocnmr.neo501 DMSO /opt/nmrdata/storage zoedg_3110 58
f1 (ppm)


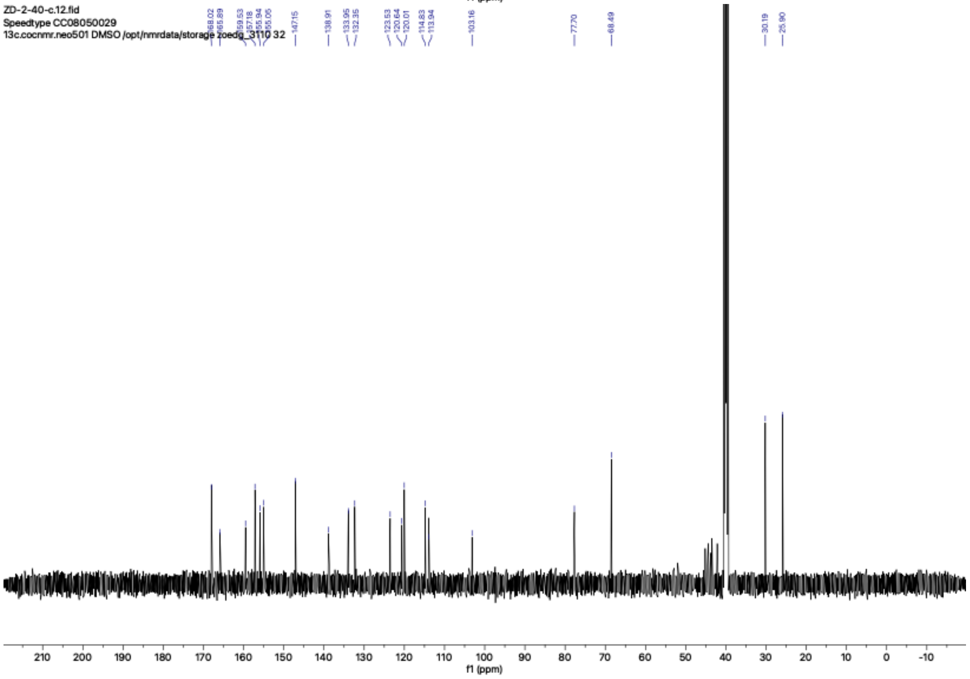

ZD-2-40-c.12.fid
Speedtype CC08050029
13c.cocnmr.neo501 DMSO /opt/nmrdata/storage zoedg_3110 32
f1 (ppm)

## ZD-1-53

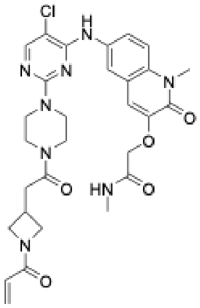

Cl
N
N
N
H
N
N
N
O
O
O
HN
O
N
O


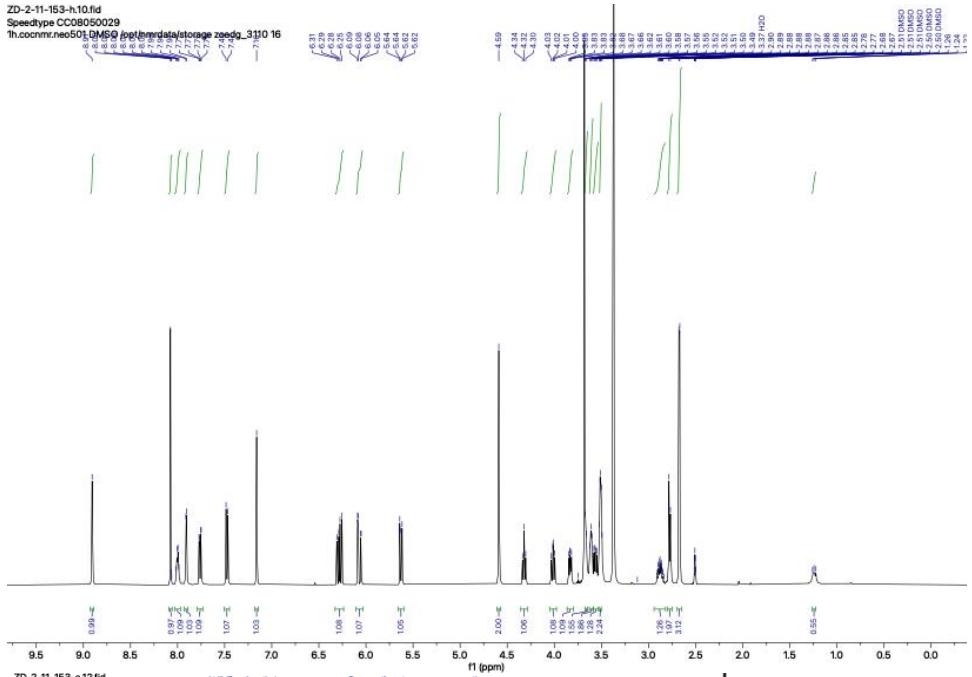

ZD-2-11-153-h.10.fid
Speedtype CC08050029
1h.cocnmr.neo501 DMSO /opt/nmrdata/storage zoedg_3110 16
f1 (ppm)


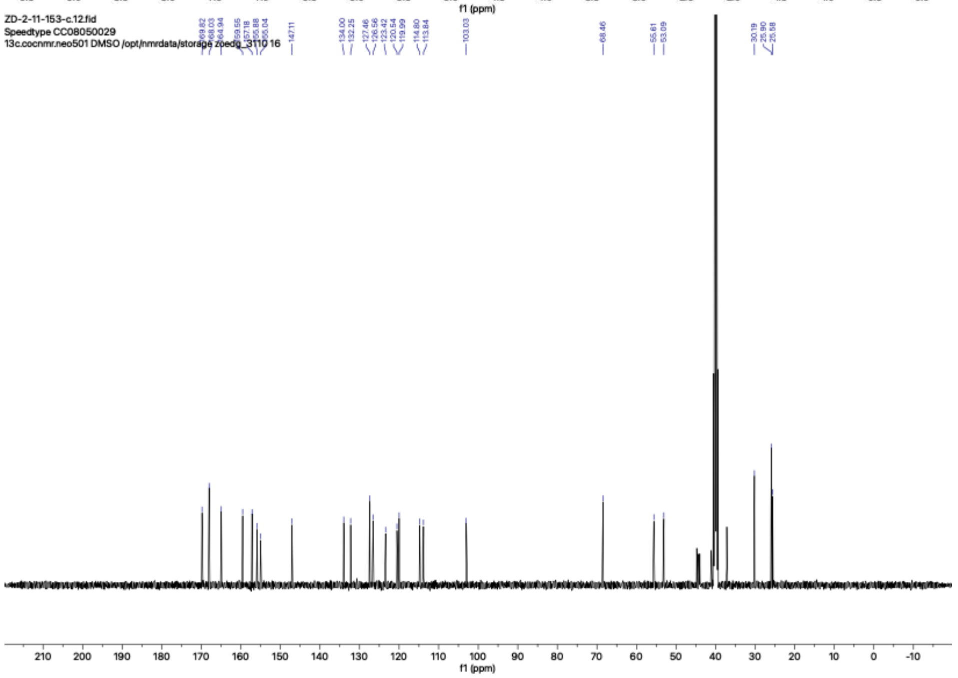

ZD-2-11-153-c.12.fid
Speedtype CC08050029
13c.cocnmr.neo501 DMSO /opt/nmrdata/storage zoedg_3110 16
f1 (ppm)

**ZD-1-55**

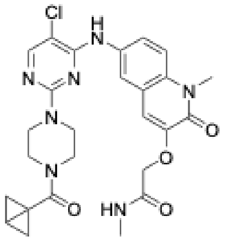

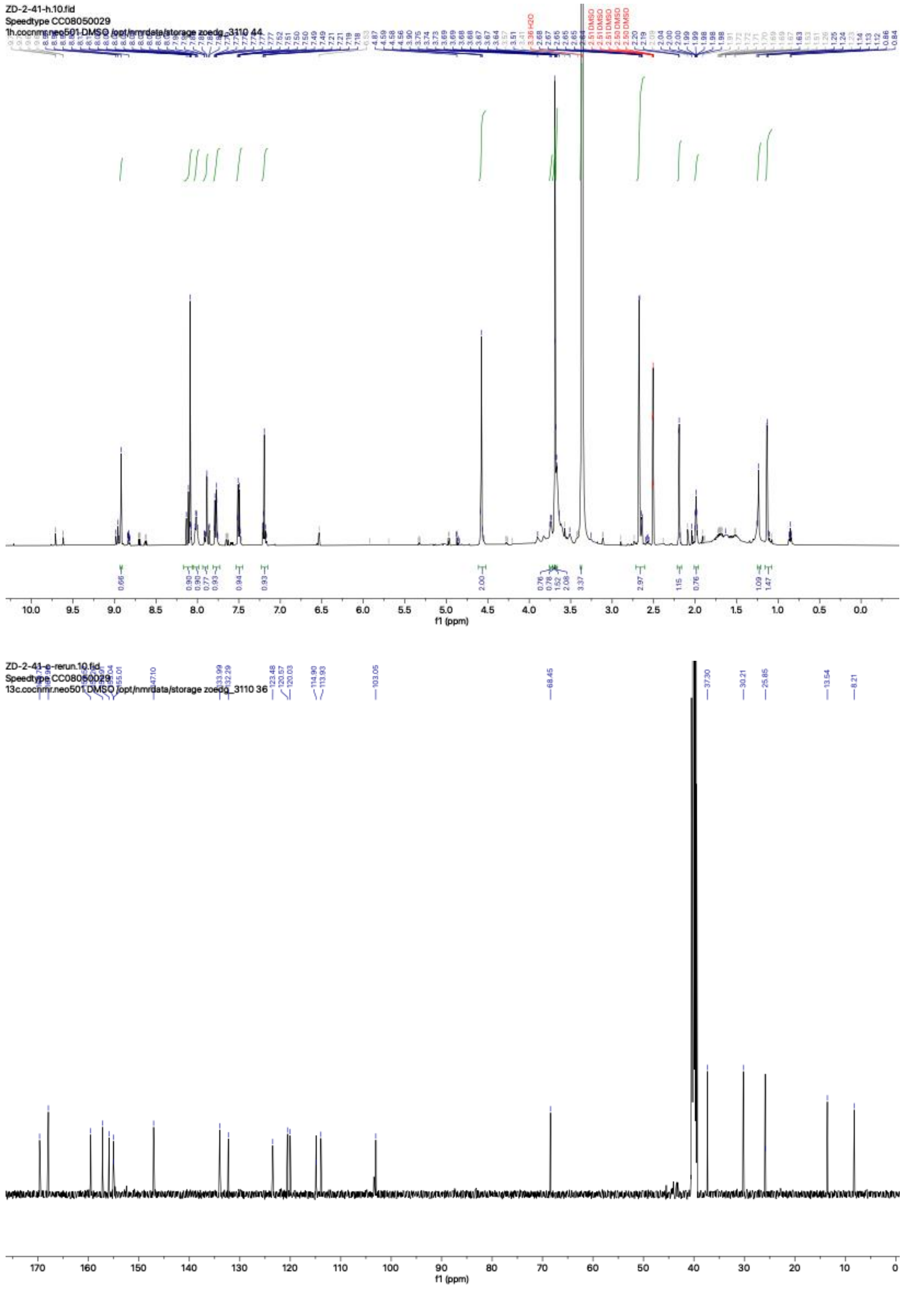

ZD-2-41-h.10.fid
Speedtype CC08050029
1h.cocnmr.neo501 DMSO /opt/nmrdata/storage zoedg_3110 44
3.36 H2O
2.51 DMSO
2.50 DMSO
f1 (ppm)
ZD-2-41-c-rerun.10.fid
Speedtype CC08050029
13c.cocnmr.neo501 DMSO /opt/nmrdata/storage zoedg_3110 36
147.10
133.99
132.29
123.48
120.57
120.03
114.90
113.93
103.05
68.45
37.30
30.21
25.85
13.54
8.21
f1 (ppm)

**4**

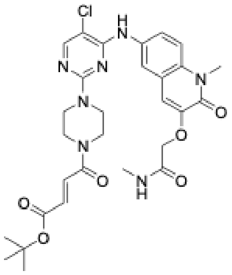

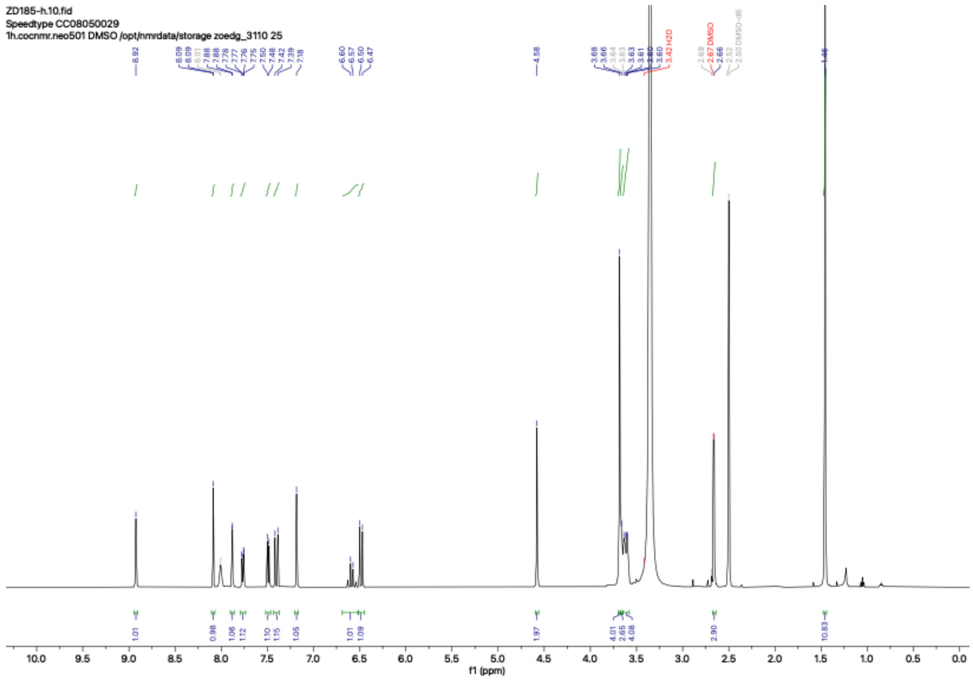
ZD185-h.10.fid
Speedtype CC08050029
1h.cocnmr.neo501 DMSO /opt/nmrdata/storage zoedg_3110 25
f1 (ppm)

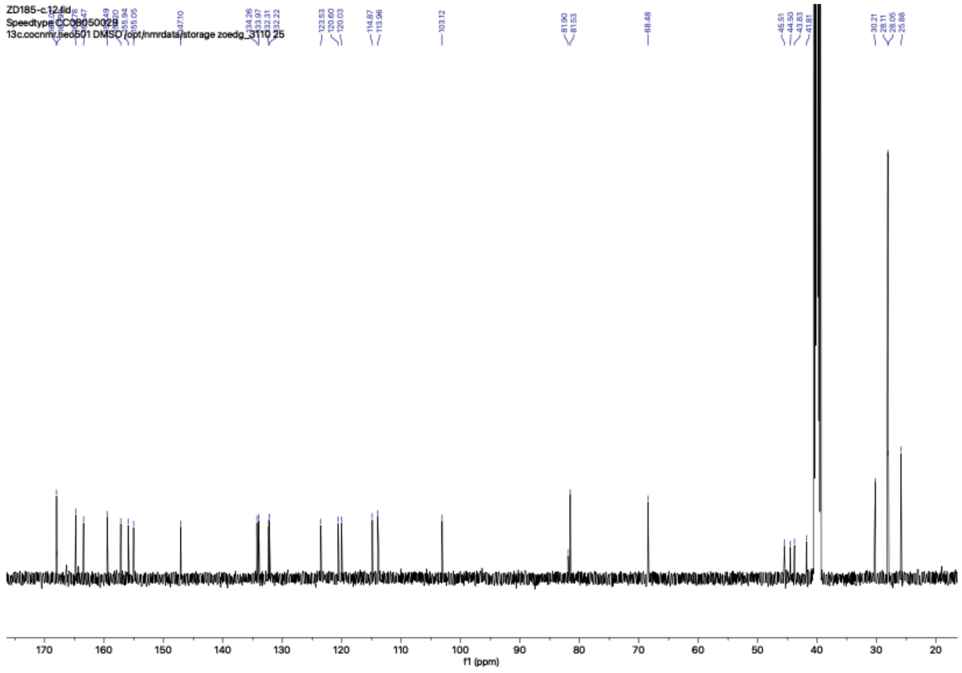
ZD185-c.12.fid
Speedtype CC08050029
13c.cocnmr.neo501 DMSO /opt/nmrdata/storage zoedg_3110 25
f1 (ppm)

**ZD-1-181**

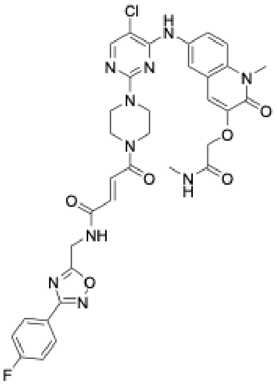

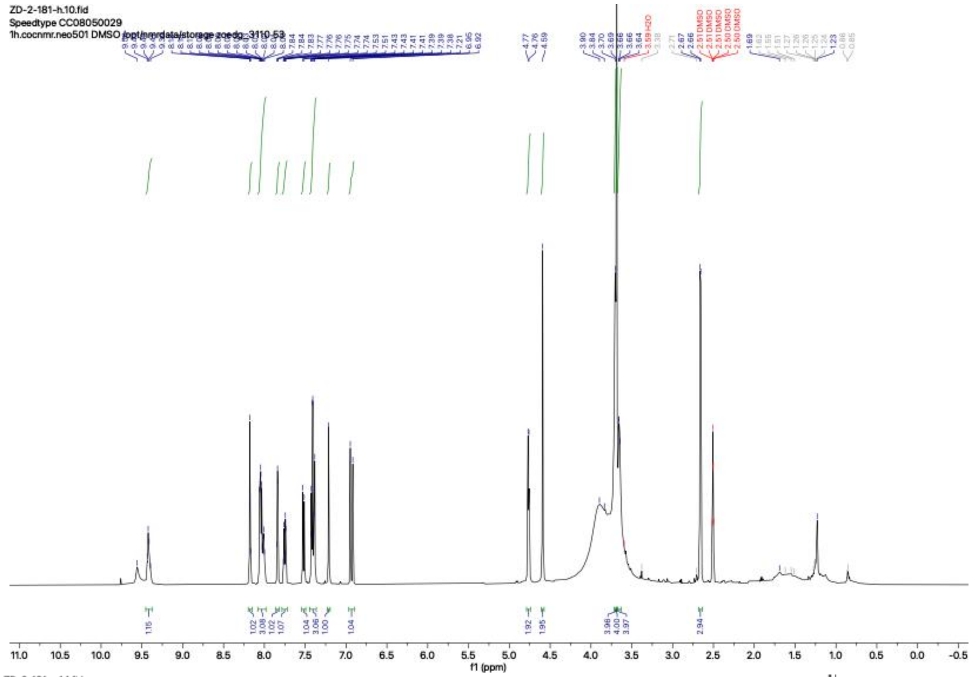
ZD-2-181-h.10.fid
Speedtype CC08050029
1h.cocnmr.neo501 DMSO /opt/nmrdata/storage zoedg_3110 53
f1 (ppm)

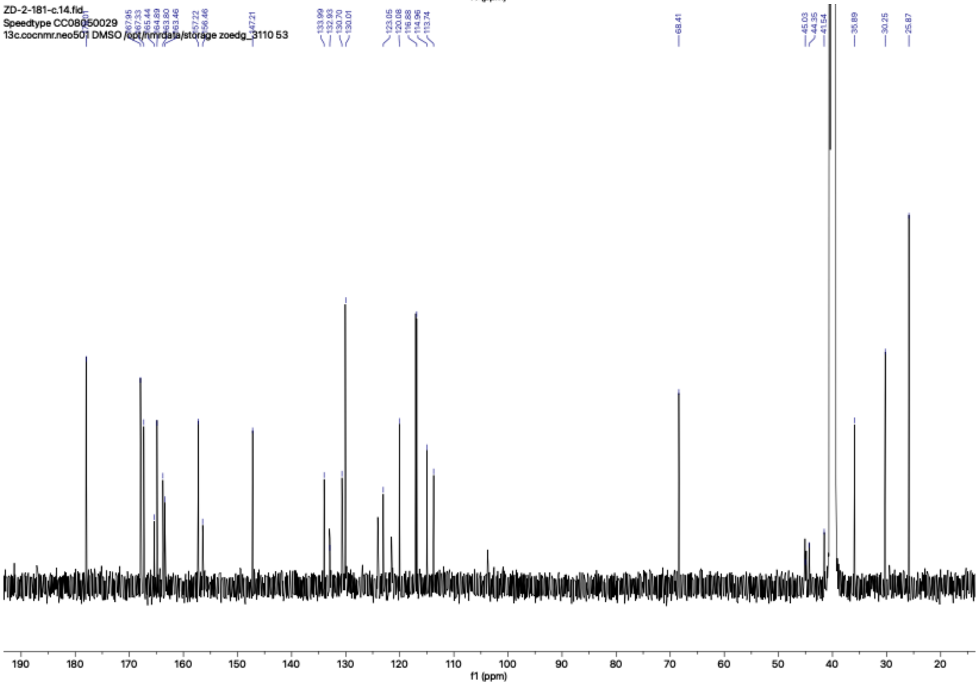
ZD-2-181-c.14.fid
Speedtype CC08050029
13c.cocnmr.neo501 DMSO /opt/nmrdata/storage zoedg_3110 53
f1 (ppm)

**ZD-1-182**

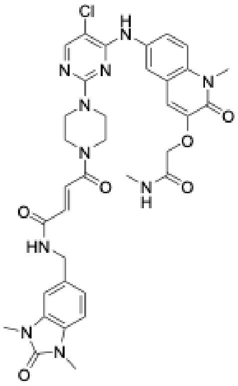

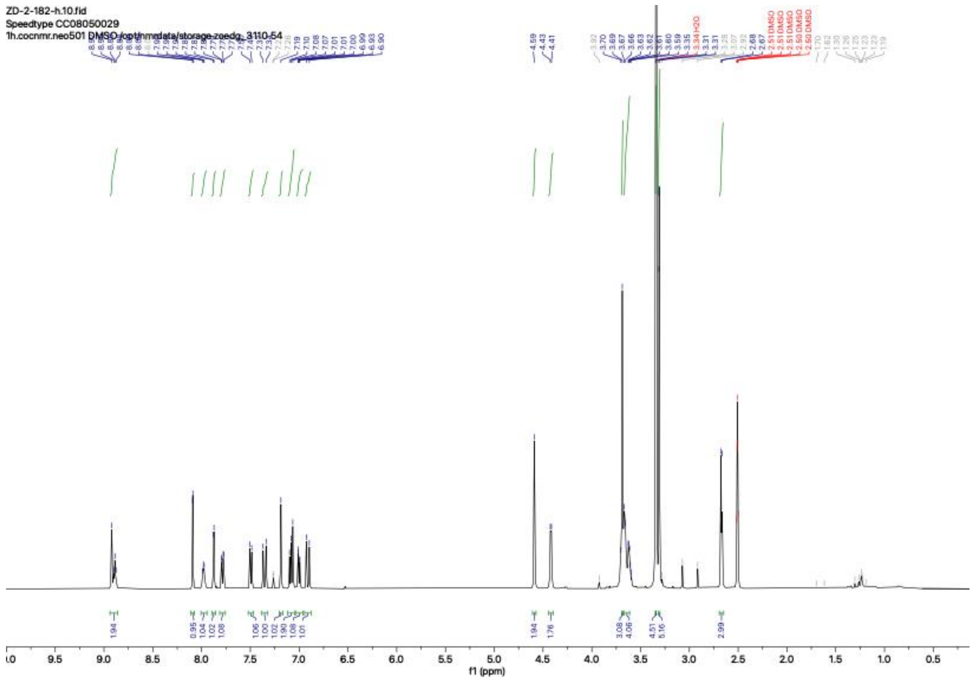
ZD-2-182-h.10.fid
Speedtype CC08050029
1h.cocnmr.neo501 DMSO /opt/nmrdata/storage zoedg_3110 54
f1 (ppm)

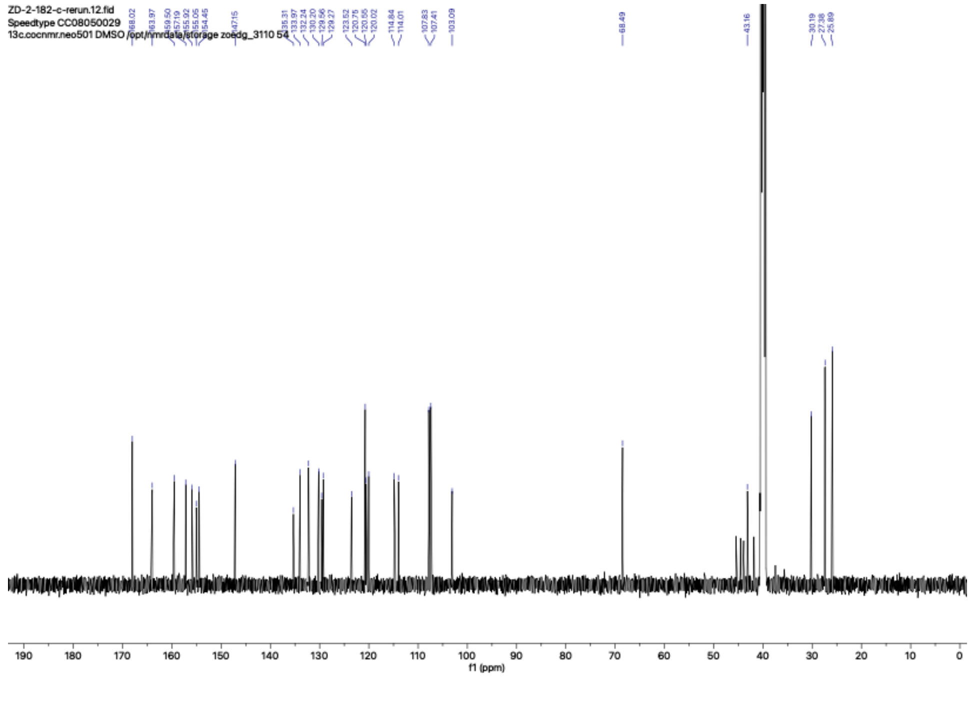
ZD-2-182-c-rerun.12.fid
Speedtype CC08050029
13c.cocnmr.neo501 DMSO /opt/nmrdata/storage zoedg_3110 54
f1 (ppm)

# ZD-1-184

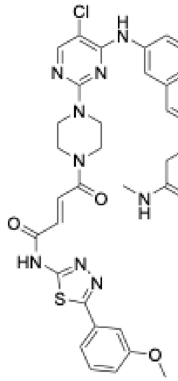

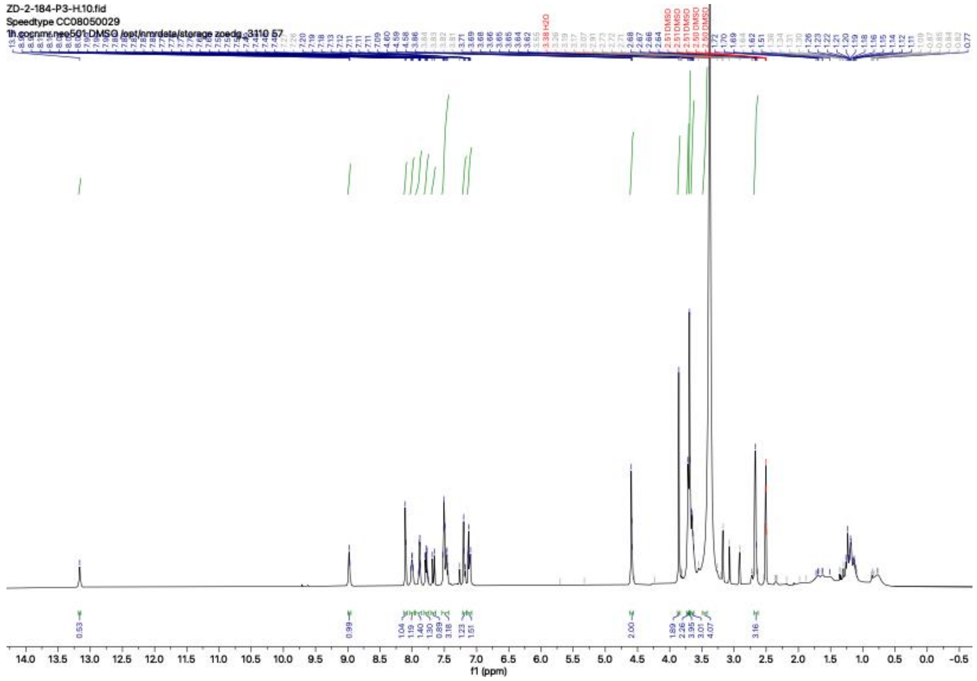

## ZD-1-186

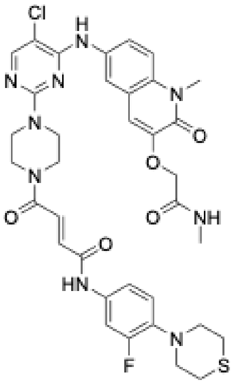
Cl
H
N
N
N
N
N
N
O
O
O
O
NH
O
HN
F
N
S

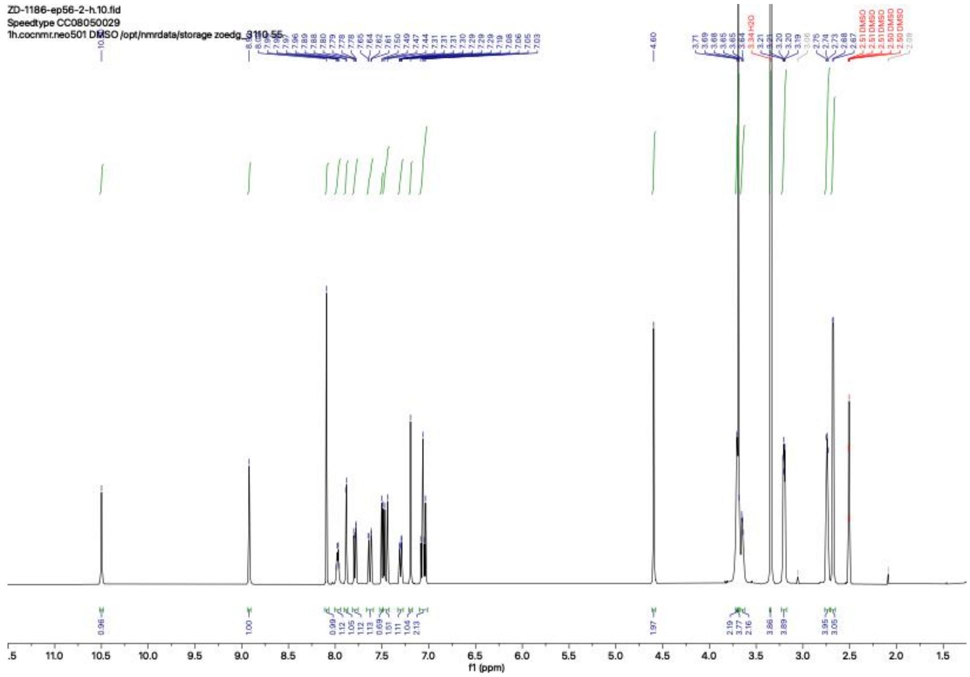
ZD-1186-ep56-2-h.10.fid
Speedtype CC08050029
1h.cocnmr.neo501 DMSO /opt/nmrdata/storage zoedg_3110 55
f1 (ppm)

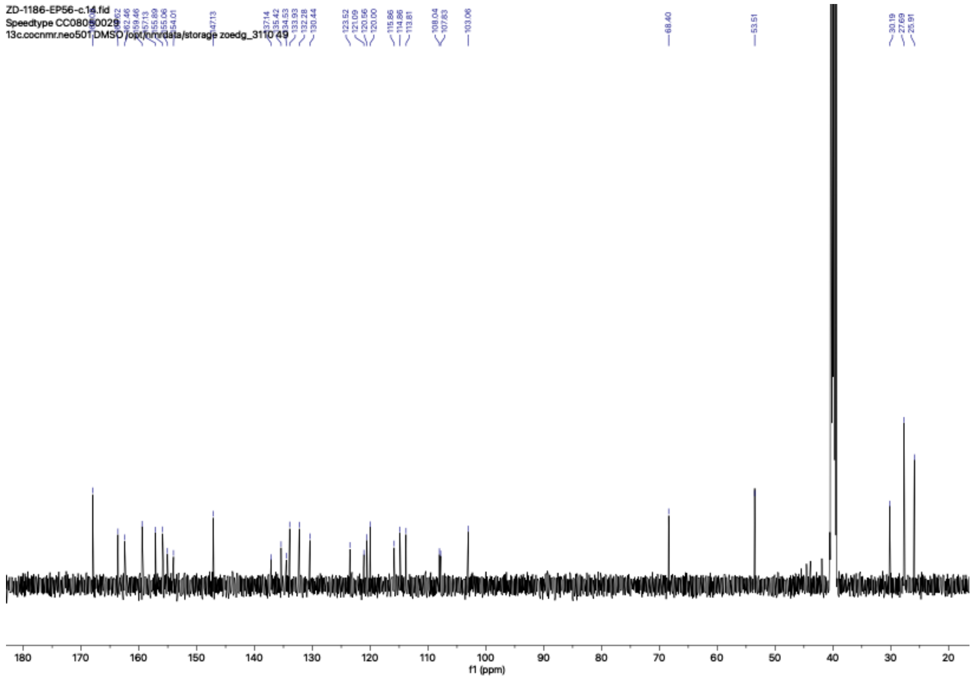
ZD-1186-EP56-c.14.fid
Speedtype CC08050029
13c.cocnmr.neo501 DMSO /opt/nmrdata/storage zoedg_3110 49
f1 (ppm)

### Compound 6

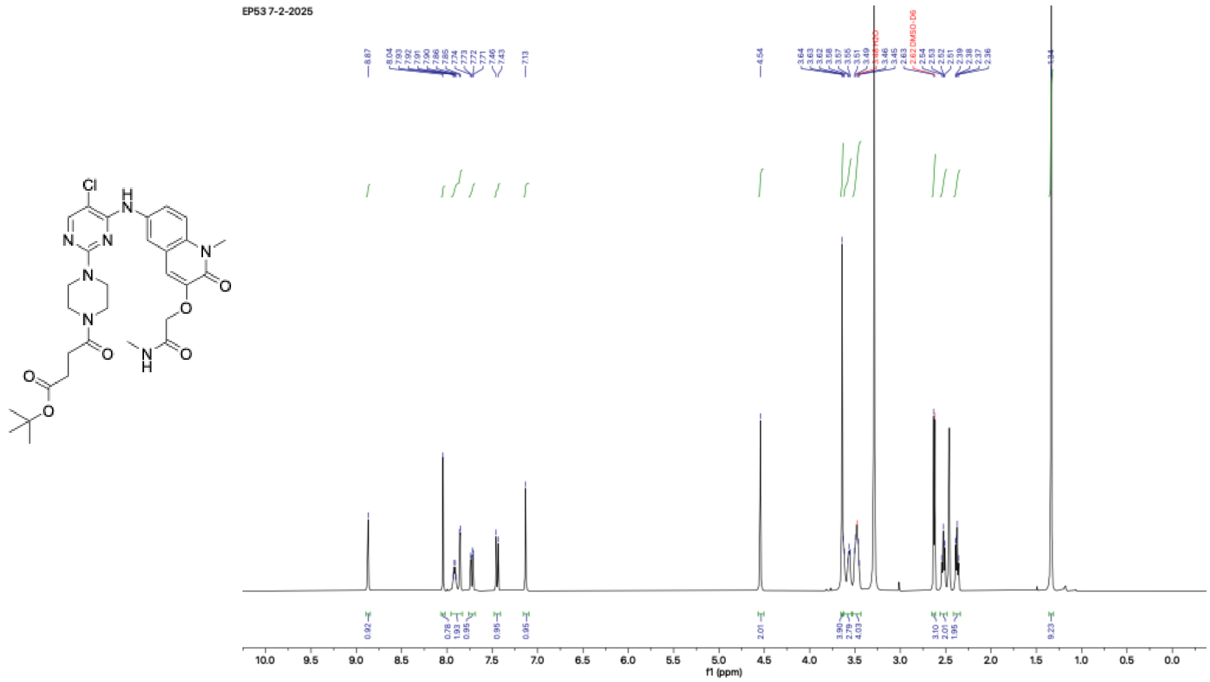


## ZD-2-2

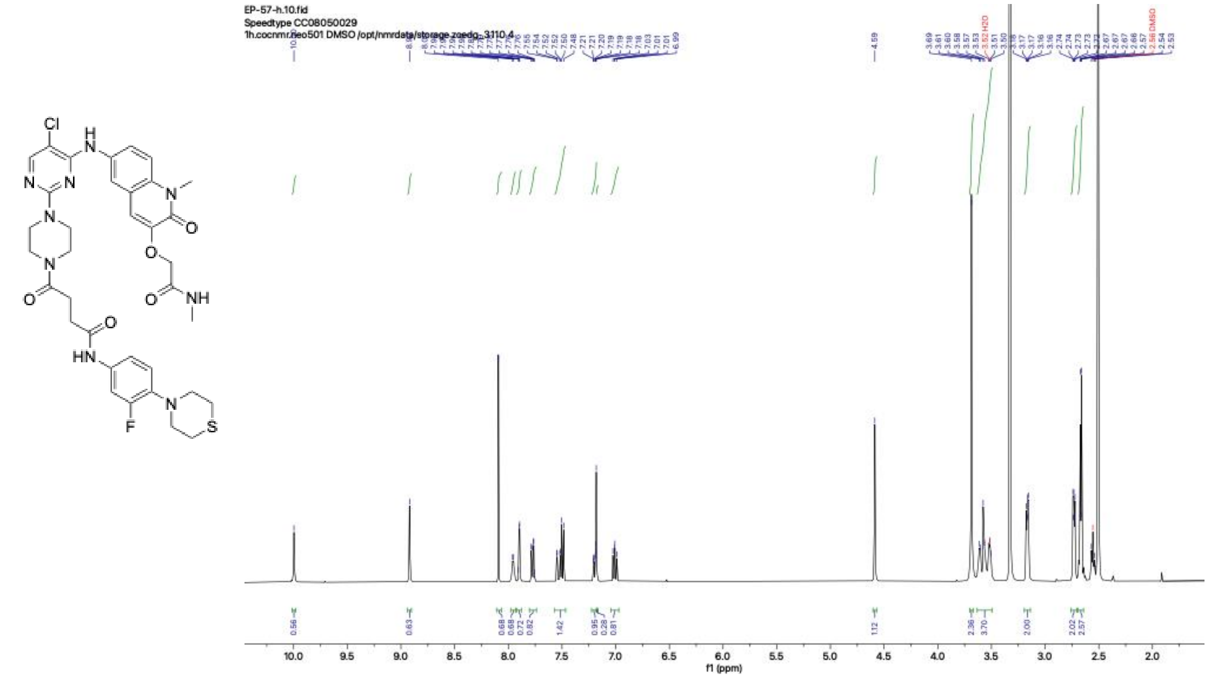
